# Single-cell chromatin profiling reveals dynamic regulatory logic and enhancer elements in brain and retina development

**DOI:** 10.64898/2026.03.16.712178

**Authors:** Jessie E. Greenslade, Hemagowri Veeravenkatasubramanian, Marisa L. Reed, Bushra Raj

**Author notes:** Corresponding author. (B.R.).

## Abstract

Cell type identity in the nervous system is encoded within cis-regulatory landscapes that integrate transcription factor activity with chromatin accessibility. However, how these regulatory programs are organized and remodeled during post-embryonic neural development remains poorly understood. We generate a temporally resolved single-cell chromatin accessibility resource of ~95,000 zebrafish brain and retina nuclei spanning larval, juvenile, and adult stages. We define a total of 213 chromatin states and uncover extensive cell type- and region-specific reorganization of chromatin landscapes across the profiled stages. By integrating scATAC-seq with transcriptomic data, we link motif accessibility to transcription factor expression and reveal distinct patterns of regulatory stability and remodeling across neural cell types and brain regions. Leveraging this resource, we systematically identify and functionally validate candidate enhancers in vivo. Focusing on the *slc1a3b* locus in radial glia, we define compact enhancer modules that act combinatorially to drive gene expression and identify candidate regulatory inputs shared with human *SLC1A3*. Together, these findings provide a framework for decoding neural cis-regulatory logic and functionally interrogating regulatory elements across the vertebrate nervous system.

## INTRODUCTION

The remarkable cellular diversity of the vertebrate nervous system emerges from spatiotemporally coordinated gene regulatory programs that guide neural progenitors through proliferation, specification, differentiation, and long-term maturation. Although single-cell transcriptomic studies have mapped neural populations at high resolution (Siletti et al. 2023; Chen et al. 2024; La Manno et al. 2021; Han et al. 2025; Yao et al. 2023; Raj et al. 2020), gene expression patterns ultimately reflect underlying cis-regulatory landscapes that govern transcription factor (TF) access to target genes. Promoters, enhancers, and related regulatory elements function combinatorially to establish and maintain cell type-specific transcriptional states. Defining these elements at single-cell resolution is therefore essential for understanding how regulatory networks shape neural diversity.

Single-cell chromatin profiling studies are defining cell type-resolved regulatory programs in the vertebrate brain and retina (Lyu et al. 2021, 2023; Zuo et al. 2024; Zu et al. 2023; Li et al. 2023). Timepoint-resolved single-cell chromatin accessibility maps further enable systems-level analysis of neural regulatory principles across both anatomical regions and developmental stages, revealing chromatin reorganization over time, variation in TF-associated regulatory logic, and cell type- and region-specific cis-regulatory elements. While it is well established that embryonic stages are characterized by highly dynamic processes at both the chromatin and transcriptional levels, a similar analysis has not been performed for post-embryonic development. Chromatin landscape dynamics across postnatal, juvenile, and adult stages, which are marked by circuit expansion and functional maturation, remain incompletely characterized, in part because of the large temporal gaps between these stages and increasing tissue complexity. Thus, it is poorly understood how the chromatin landscapes of cell types and regions are remodeled during post-embryonic neurogenesis and which TF combinations and regulatory programs define these populations over time.

The compact size and accessibility of the zebrafish nervous system provide a practical platform for whole-brain and whole-retina single-cell chromatin profiling across post-embryonic stages. Prior zebrafish single-cell ATAC sequencing (scATAC-seq) studies have largely centered on early timepoints and whole-embryo or whole-animal preparations without enrichment of neural populations (Macho Rendón et al. 2026; Kim et al. 2024; Sun et al. 2024; Xu et al. 2024; McGarvey et al. 2022; Liu et al. 2024). Here, we generate a single-cell chromatin accessibility resource of the zebrafish brain and retina spanning larval, juvenile, and adult stages, providing detailed characterization of cis-regulatory dynamics across post-embryonic periods of neural development. The annotated resource and underlying chromatin tracks are available to the community for interaction and exploration through the UCSC genome browser. By integrating scATAC-seq with previously published transcriptomic datasets, we define cell type- and region-specific TF regulatory landscapes and chart their temporal stability across post-embryonic development. Taking advantage of the optical transparency of larval zebrafish, we leverage this resource to systematically identify and functionally validate cis-regulatory elements in vivo with reporter constructs. Focusing on the *slc1a3b* gene, we resolve its enhancer architecture through fine-scale functional mapping and mutation analyses. Together, these studies provide a framework for linking post-embryonic chromatin dynamics to cell type- and region-specific regulatory programs and for functionally interrogating the cis-regulatory elements that shape neural gene expression.

## RESULTS

### Mapping chromatin accessibility in the larval, juvenile and adult zebrafish brain and retina

We profiled chromatin accessibility across three stages of zebrafish brain and retina development: 3 days post-fertilization (dpf), shortly after the onset of secondary neurogenesis when larval neural tissues rapidly expand; 21 dpf, a juvenile stage during continued neural growth; and 5 months post-fertilization (5 mpf), representing a mature adult brain and retina (Fig. 1A, 1B). To enrich for brain and retinal populations, we dissected whole heads at 3 dpf, and isolated brains and eyes at 21 dpf and adult stages. Nuclei were processed using the 10x Chromium scATAC-seq platform. To increase adult coverage, we incorporated previously published scATAC-seq profiles from ~20,000 adult brain nuclei (Yang et al. 2020). After quality filtering, we retained 94,976 high-quality nuclei across the three stages with a median of 10,110 fragments per nucleus.

**Fig 1.**
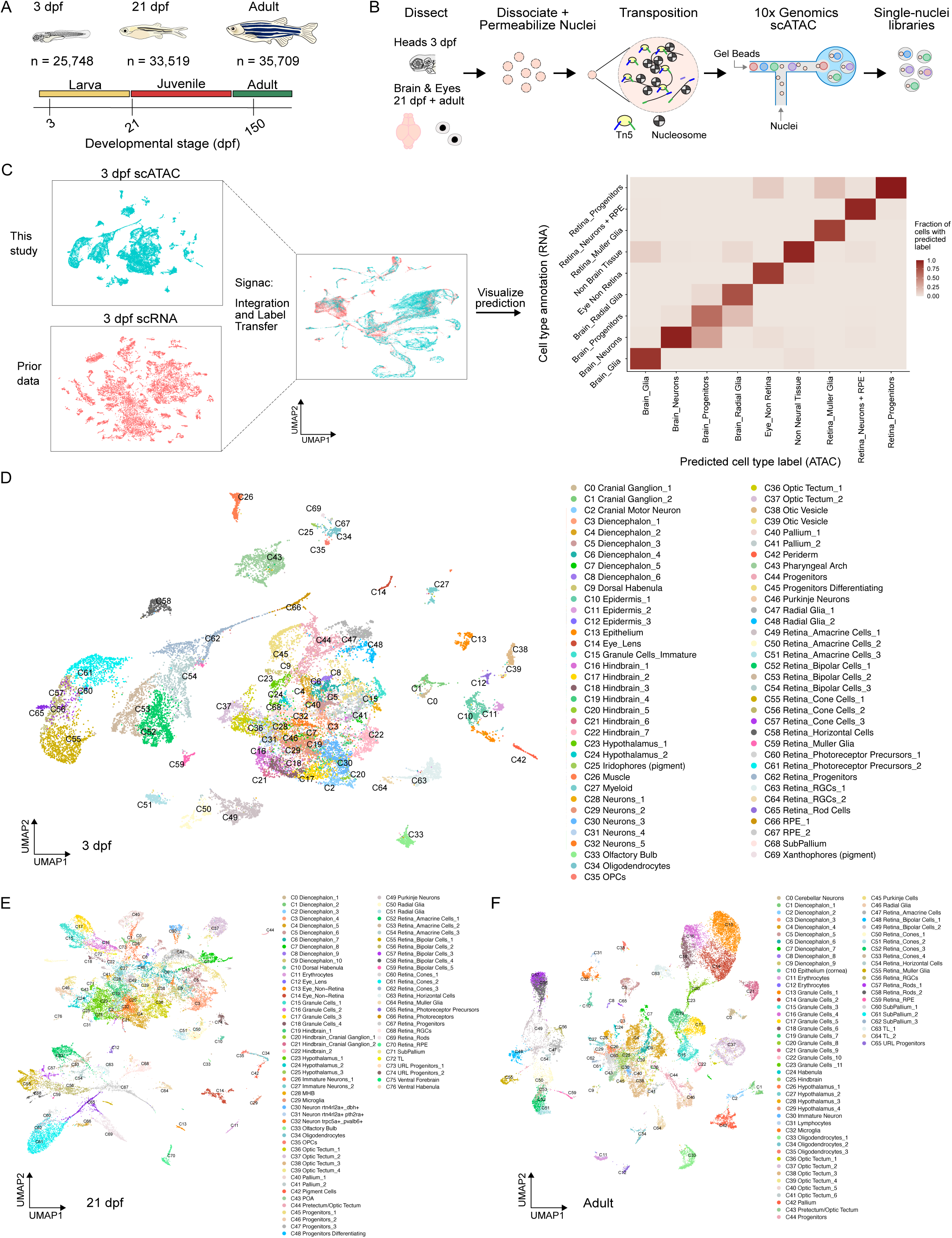
Single-cell chromatin accessibility datasets of the zebrafish brain and retina. (A) Overview of developmental stages profiled, and the number (n) of brain and eye nuclei analyzed by scATAC-seq. dpf, days post-fertilization. (B) Experimental overview of scATAC-seq workflow. (C) scATAC-seq cell type annotation strategy. Right, confusion matrix showing agreement between scRNA-seq broad cell type labels and predicted scATAC-seq cell type assignments following label transfer. 3 dpf data is shown as an example; scRNA-seq data is from (Raj et al. 2020). (D-F) UMAP plots of scATAC-seq profiles at (D) 3 dpf, (E) 21 dpf, and (F) adult stages, colored by annotated cell type.

We clustered each dataset independently and integrated the scATAC-seq profiles with our previously published scRNA-seq datasets (Raj et al. 2020; Siniscalco et al. 2024), using the Signac R package (Stuart et al. 2021). scRNA-derived cell type labels were projected onto scATAC clusters by label transfer (Fig. 1C). Cell type assignments were manually reviewed and confirmed by the enrichment of accessible chromatin peaks around canonical marker genes (Supplemental Fig. S1, Supplemental Table S1). Neuronal cell type annotations include their predicted regional localization (e.g. Diencephalon) or named cell type (e.g. Bipolar Cell) derived from accessibility around spatially restricted TFs and known marker genes (Fig. 1D-F). Subtypes were further resolved by cluster-specific accessibility patterns in addition to shared regional or cell type signatures (Supplemental Fig. S2, Supplemental Table S2). Non-neural clusters, such as pigment cells and cell types from the pharyngeal arch, muscle, periderm, and epidermis were detected primarily at 3 dpf due to the inclusion of whole heads and were excluded from downstream analyses. We classified 70 clusters at 3 dpf, 77 clusters at 21 dpf, and 66 clusters in the adult, resulting in a total of 213 chromatin states. Collectively, our datasets captured major neural and retinal populations, providing a high-resolution resource of chromatin accessibility across post-embryonic zebrafish brain and retina development. They are available for viewing on the interactive UCSC Cell Browser website: https://zebrafish-neural-atac.cells.ucsc.edu.

### scATAC-seq captures shifts in cell populations from larval to adult stage

As developmental changes in chromatin state can arise not only from intrinsic regulatory programs but also from shifts in cellular composition, we first quantified how major cell populations change across time. In the retina, retinal progenitor cells and photoreceptor precursors from larval and juvenile stages were largely absent in the adult (Fig. 2A). At 3 dpf, 61.3% of the retina comprised inner retinal neurons (amacrine, bipolar, horizontal, and retinal ganglion cells [RGCs]), and 26.8% comprised photoreceptor cells (rods, cones, and photoreceptor precursors). By adulthood, this balance shifted: photoreceptors expanded to 57.8% of the retina, whereas inner retinal neurons decreased to 34.1%. These trends align with established patterns of zebrafish retinal neurogenesis in which RGCs, horizontal, amacrine, and cone cells arise early and bipolar cells, rods, and Müller glia emerge later (Lyu et al. 2023; Hoang et al. 2020). Consistent with this trajectory, cones were already abundant by 3 dpf and increased modestly thereafter, while rods showed the most pronounced rise, increasing from 1.2% at 3 dpf to 29.4% in the adult (Fig. 2B). In the brain, granule cells exhibited the largest expansion of any lineage, increasing from 1.6% at 3 dpf to 37.4% in the adult, consistent with their known proliferative burst during the juvenile-to-adult period (Kaslin et al. 2013). Brain progenitor populations declined substantially (14.3% to 1.8%), while radial glia decreased more modestly (5.5% to 3.2%) from 3 dpf to adulthood. Together, these results demonstrate that our dataset captures expected post-embryonic developmental shifts in retinal and brain composition, providing a strong foundation for interpreting cell type- and region-specific chromatin accessibility changes.

**Fig 2.**
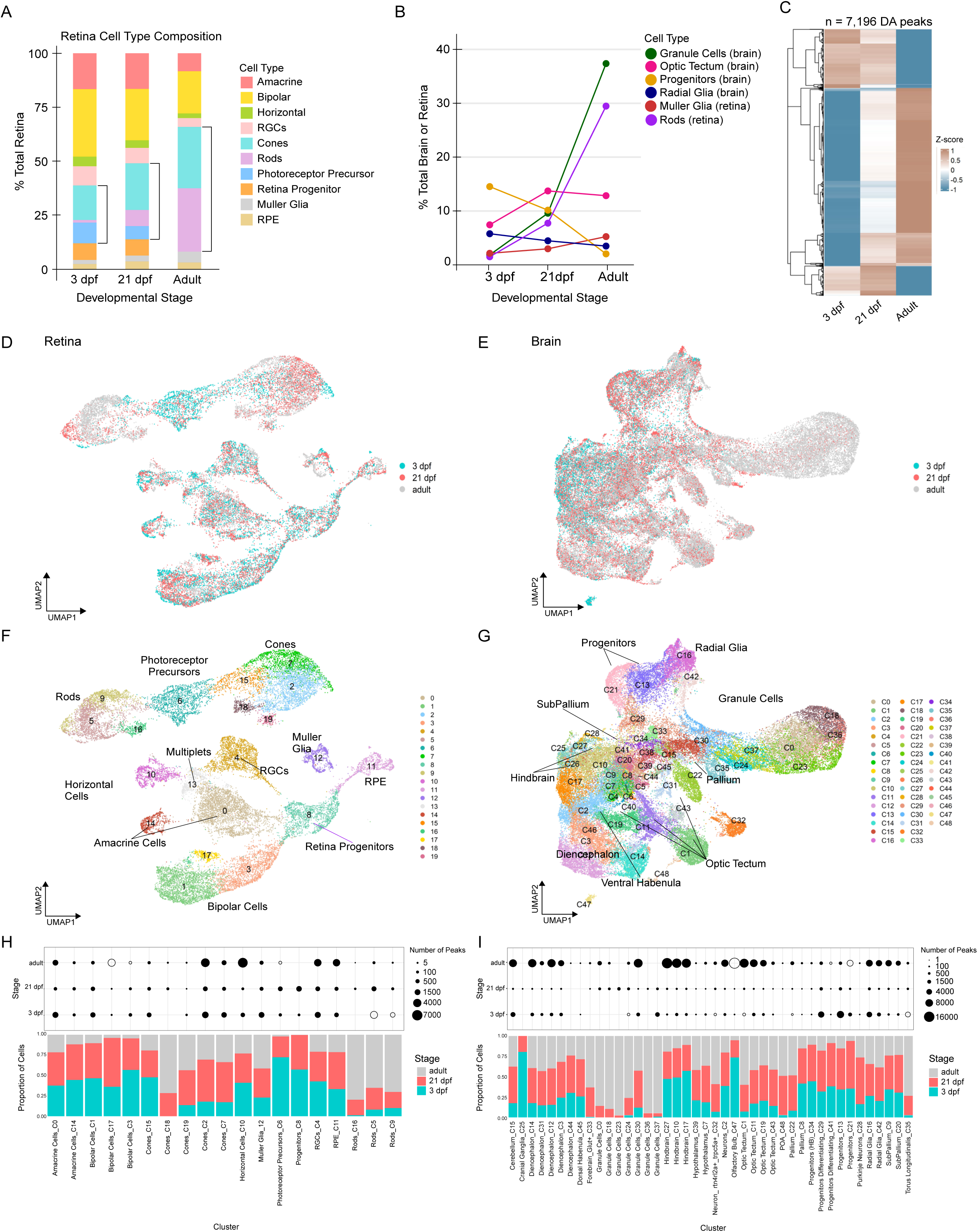
Cell type-resolved chromatin dynamics across zebrafish brain and retinal development. (A) Proportions of major retinal cell populations at each profiled stage. (B) Changes in the proportions of selected brain and retinal populations across stages. (C) Heatmap showing Z-scores of 7,196 differentially accessible (DA) peaks (log2 fold change >1, adjusted p-value < 0.05) enriched at 3 dpf, 21 dpf, and adult stages. (D-E) Harmony integrated UMAP embeddings of scATAC-seq data from the (D) retina or (E) brain showing cells colored by timepoint. (F-G) Integrated UMAP embeddings of scATAC-seq profiles of (F) retinal cells and (G) brain cells colored by annotated cell type. (H-I) Cell type-resolved DA peaks across timepoints. Dot plots show the number of stage-enriched DA peaks within each (H) retina and (I) brain cluster. Dot size reflects the number of DA peaks. Open circles indicate timepoints contributing fewer than 10% cells to a given cluster. Bar plots indicate the relative proportion of cells from each timepoint within that cluster.

### Post-embryonic cell-type and region-specific reorganization of chromatin landscapes

We next examined stage-dependent changes in the overall brain and retinal chromatin accessibility landscapes using dataset-wide comparisons across the profiled post-embryonic periods. Out of 383,101 total consensus peaks in the three datasets, we identified only 7,196 unique peaks (log2 fold change > 1), representing 1.9% of consensus peaks, with stage-specific differential accessibility (DA) in global comparisons (Fig. 2C). These regions formed distinct early- and late-accessible modules, with 21 dpf typically exhibiting intermediate accessibility, indicating progressive chromatin remodeling across the profiled stages.

To examine chromatin accessibility dynamics within cell types and regions, we separately integrated and clustered retinal and brain cells across the three timepoints (Fig. 2D-G, Supplemental Table S3). We then performed pairwise DA analyses across timepoints within each cluster to identify stage-enriched regulatory elements. We identified 35,674 unique DA peaks (9.31% of consensus peaks) in the retina and 64,898 unique DA peaks (16.94% of consensus peaks) in the brain (Fig. 2H, 2I, Supplemental Fig. S3A, S3B). Thus cluster-specific analysis revealed additional chromatin-level changes that were masked in global analysis at the dataset-wide level. DA peak abundance was not significantly associated with cluster size in the retina (Spearman’s ρ = 0.222, p = 0.106, adj. R² = 0.023) and showed only a weak association in the brain (ρ = 0.266, p = 0.005, adj. R² = 0.071), indicating that cell number is not a major determinant of DA patterns (Supplemental Table S4). The magnitude of change varied markedly across populations (Supplemental Table S5). For example, in the retina, RGCs (C4) were characterized by 2,250 larval-enriched peaks, 1,225 juvenile-enriched peaks, and 3,818 adult-enriched peaks. Similar trends were observed in the cone photoreceptor clusters C2 and C7. We also noted several clusters with adult-enriched DA profiles. For example, horizontal cells (C10) had the largest number of adult-enriched DA peaks in the retina, indicating large regulatory reorganization between the juvenile and adult stages. In the brain, subtypes of neurons in the diencephalon, hindbrain, optic tectum, and subpallium had enrichment of DA peaks in the adult stage. Radial glia displayed a bimodal pattern, with unique peak sets at both larval and adult stages and a relative depletion at the juvenile stage. Rod photoreceptors (C16 and C9), granule cells (excluding C30), dorsal habenula (C45), and cranial ganglia (C25) had relatively few DA peaks at any of the profiled stages, suggesting that chromatin identity in these cell types is established early and maintained. Remarkably, despite the pronounced expansion of rod photoreceptors and granule cells in adults, they did not show a corresponding increase in adult-enriched DA peaks.

To investigate the biological programs underlying these dynamics, we identified candidate target genes associated with DA peaks in each cluster. We calculated peak-gene correlations and identified genes whose accessibility was correlated with DA peaks (p-value < 0.05, |score| ≥ 0.05, distance to TSS within 500 kb). In the retina, genes linked to larval DA peaks include developmental signaling genes such as *notch1a* and developmental TFs such as *pax6a*, *pax6b*, and *otx5.* In contrast, genes linked to adult-enriched DA peaks include *abca4*, *guca1d*, and *gnat2* phototransduction genes in photoreceptors and *syt1a* and *stx3b* synaptic genes in amacrine cells. Gene Ontology analysis confirmed that larval-enriched DA peaks in brain and retina were associated with developmental processes including central nervous system development, axon development, and cell adhesion; while, adult-enriched peaks were linked to mature neuronal functions including visual perception (retina), synaptic signaling, calcium ion transport, and circadian regulation (Supplemental Fig. 3C; Supplemental Table S6).

### Cell type- and region-resolved transcription factor motif analysis

To characterize cell type- and region-associated regulatory programs, we applied chromVAR (Schep et al. 2017) to compute per-cell TF motif deviation scores, using a zebrafish-specific TF motif database (SwissRegulon). Enrichment of motif-containing accessible sites serves as a proxy for inferred TF regulatory potential. We focused initially on the 3 dpf scATAC-seq dataset and identified motif accessibility patterns across 57 brain and retinal clusters (excluding non-neural clusters). Hierarchical clustering of the top 25 accessible motifs per cluster revealed regulatory signatures corresponding to distinct neural and retinal lineages (Fig. 3A, Supplemental Table S7). For example, motifs associated with photoreceptor specification, including *crx* and *otx5* (co-orthologs of mammalian Crx), showed elevated accessibility in photoreceptor populations, whereas *pou4f3* motif accessibility was enriched in the RGC cluster (Fig. 3B). SoxB1 family motif was enriched in accessible chromatin regions within neural progenitor and radial glia populations, consistent with stem- and progenitor-associated regulatory states. Motif accessibility for the regionally restricted factors *hoxa2b* and *lef1* was enriched in hindbrain- and diencephalon-associated clusters, respectively. We observed similar lineage-specific patterns of motif accessibility at 21 dpf and in adults (Supplemental Fig. S4, S5, Supplemental Table S7).

**Fig 3.**
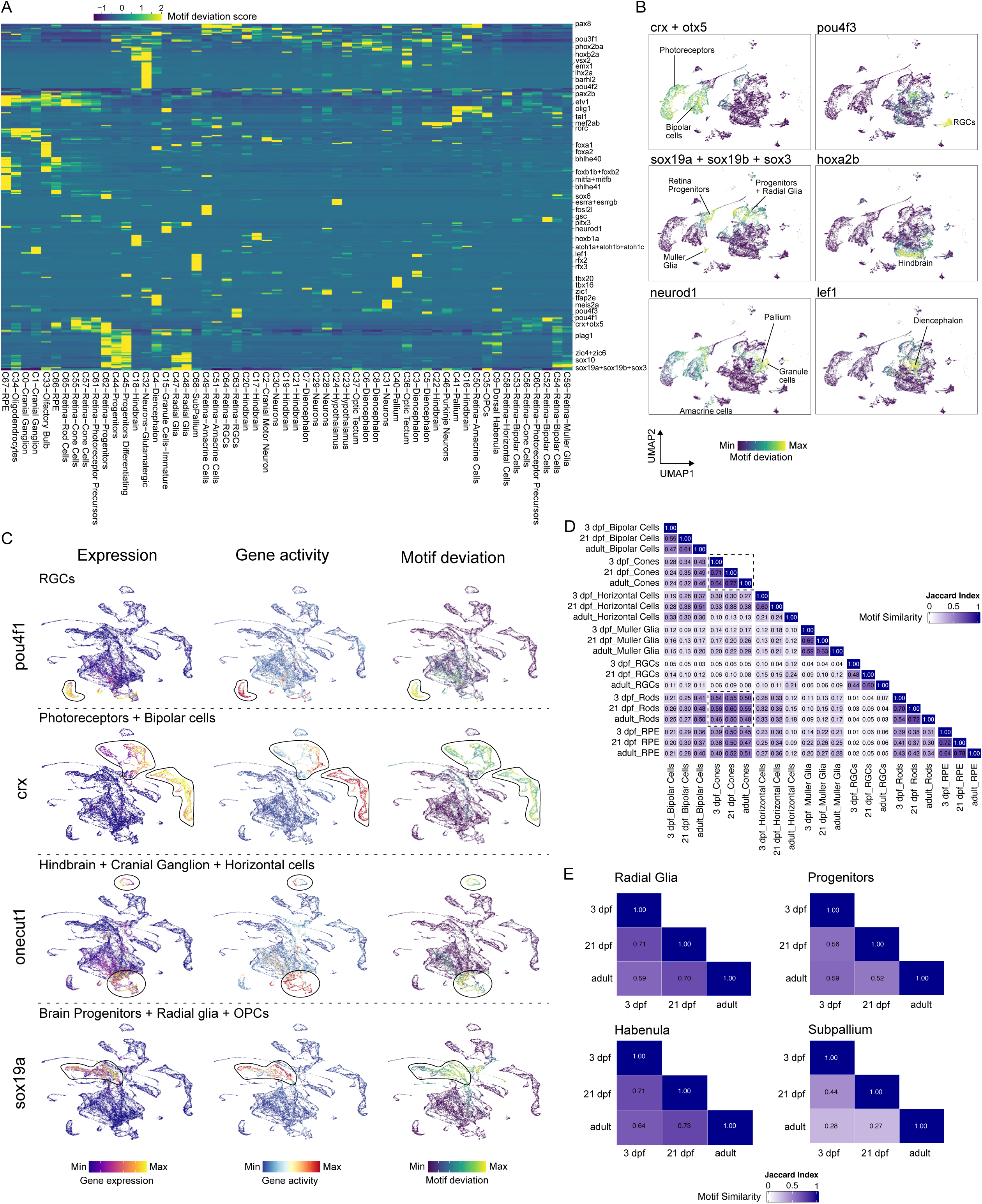
Transcription factor motif accessibility reveals cell type- and region-specific regulatory programs. (A) Heatmap of chromVAR TF motif deviation scores across 57 clusters at 3 dpf. (B) UMAP projections of 3 dpf scATAC-seq data colored by chromVAR motif deviation scores for selected TFs. TFs with similar motifs are grouped together and separated by the plus sign. (C) UMAP projections of the integrated 3 dpf scATAC-scRNA dataset colored by motif accessibility, gene activity, and gene expression for selected TFs. (D-E) Similarity of aggregate chromVAR-derived motif accessibility profiles within cell types and regions across timepoints. Heatmap shows Jaccard index overlap. Dashed boxes indicate rods and cones.

To further support TF regulatory activity inferred from motif accessibility, we compared motif enrichment with corresponding TF gene expression and gene activity (accessibility around the gene body) in the integrated 3 dpf scATAC-scRNA dataset (Supplemental Fig. S6A). We identified TFs for which cluster-enriched motif accessibility coincided with increased gene activity and expression in the same populations (Fig. 3C, Supplemental Fig. S6B). For example, *pou4f1* is highly expressed in RGCs, its locus shows increased gene activity in the same cells, and its cognate motif is preferentially accessible in the same population. Concordance across these three measures links TF availability with accessibility of both the TF locus and its cognate regulatory sequence, supporting a role for these TFs in establishing or maintaining cell type and region-specific regulatory chromatin states.

In retinal cell types, multiple members of TF families are known to be co-expressed as part of core regulatory programs that define the major retinal cell classes. We therefore asked whether these established lineage-associated TF signatures could be recovered from motif accessibility profiles. Indeed, motifs corresponding to known retinal TF families showed cell type-enriched accessibility patterns that were largely maintained across the profiled stages (Supplemental Fig. S7A). These include motifs corresponding to the *otx* family in photoreceptor lineages, *tfap2* family in amacrine cells, *pou* family in RGCs, *onecut* family in horizontal cells, and *sox* and *tead* families in Müller glia. Thus, our analyses recapitulated known retinal lineage-defining TF signatures.

We next asked whether these lineage-specific patterns extend beyond selected TF families to the broader regulatory motif repertoire of each retinal cell type. We compared aggregate chromVAR-derived motif accessibility profiles between cell types and temporal stages. Most retinal cell types had higher within-cell type similarity across timepoints than between cell types (Fig. 3D), indicating persistence of broader TF regulatory programs. Aggregate motif profiles also captured relationships between related retinal lineages; for example, rods and cones were more similar to one another than to other retinal cell classes, consistent with their shared photoreceptor regulatory identity.

Brain regions are defined by combinatorial TF activity that establishes and maintains regional identity, but how these TF regulatory programs change during post-embryonic neural development remains poorly understood. To assess this, we compared chromVAR-derived motif profiles for corresponding brain regions across the profiled timepoints. The subpallium, pallium, diencephalon, optic tectum, and hindbrain regions showed relatively low overlap in accessible TF motifs across timepoints, suggesting substantial remodeling of their regulatory programs during post-embryonic development (Fig. 3E, Supplemental Fig. S7B, Supplemental Table S8). For example, motifs associated with GABAergic subpallial identity, including members of the *dlx*, *lhx*, and *mef2* families, were shared between larval and juvenile stages but absent in the adult. These observations were consistent with stage-dependent remodeling of the subpallial regulatory landscape. In contrast, the habenula exhibited relatively high motif-profile similarity across stages, with *pou4f1*, *nr4a2a*, and *nr4a2b* motifs retained across timepoints, suggesting greater temporal stability of its regulatory program.

Finally, we extended these analyses to defined brain cell types. Among progenitor-associated cell types, radial glia showed greater motif similarity across all three timepoints than brain progenitors (Fig. 3E). This may reflect maintenance of a more persistent core regulatory program in radial glia, consistent with their long-term neural stem cell-like identity in the zebrafish brain, whereas brain progenitors undergo greater stage-dependent remodeling of their regulatory states. Cerebellar granule cells and Purkinje cells showed low motif overlap when comparing 3 dpf to the other stages, but greater similarity between juvenile and adult stages (Supplemental Fig. S7B), suggesting that their regulatory programs undergo early remodeling before becoming more stable later in development. Collectively, these analyses reveal that TF-associated regulatory programs differ substantially in their temporal stability across neural cell types and brain regions, providing a framework for linking chromatin accessibility dynamics to cell type- and region-specific regulatory logic during neural development.

### Identification of candidate enhancer elements

Our resource also enables discovery of cis-regulatory elements (CREs) in the brain and retina. Genomic annotation of the 383,101 consensus peaks showed that the majority mapped to non-coding DNA, with 65% and 25.52% located in intergenic and intronic regions, respectively (Fig. 4A). To identify candidate CREs with cell type- or region-restricted activity, we developed a bioinformatic pipeline that integrates differential chromatin accessibility with differentially expressed genes from scRNA-seq (Supplemental Fig. S8). Candidate CREs were prioritized based on strong differential accessibility, restricted accessibility across cell types or regions, and proximity to differentially expressed genes in the corresponding scRNA-seq populations. Since intergenic peaks comprised the largest fraction, we focused on these regions for functional validation.

**Fig 4.**
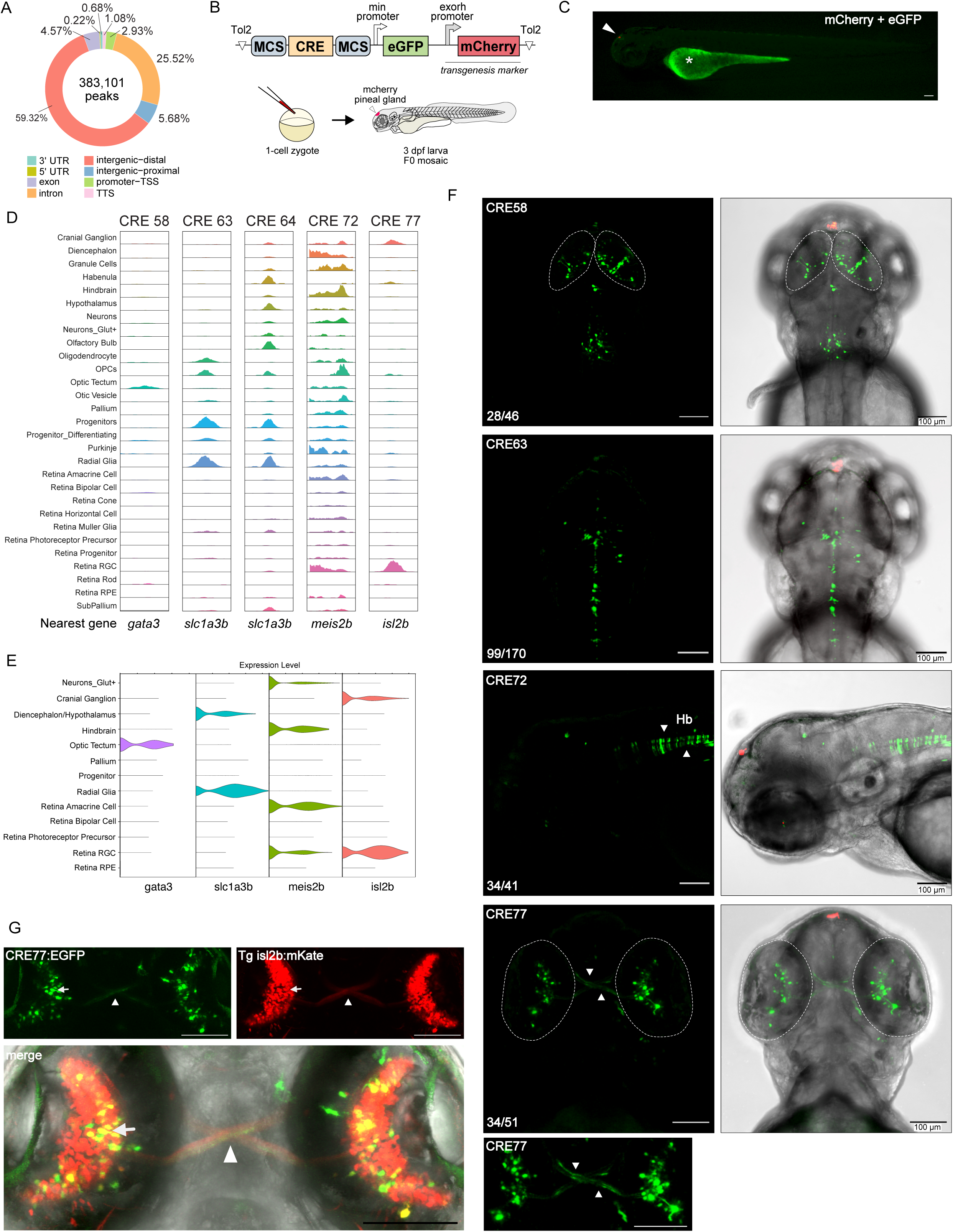
In vivo validation of candidate cis-regulatory elements. (A) Genomic annotation of consensus chromatin accessibility peaks. TSS, transcription start site. TTS, transcription termination site. (B) Schematic of the enhancer reporter assay. Reporters were injected into 1-cell stage zebrafish embryos. Reporter activities were assessed at 3 dpf with mCherry serving as a transgenesis marker. MCS, multiple cloning site. (C) Fluorescence image of 3 dpf zebrafish injected with an empty reporter plasmid. No eGFP is detected while mCherry (arrowhead) marks successful transgenesis. Asterisk, autofluorescence in yolk. (D) Aggregate chromatin accessibility profiles of selected candidate CREs at 3 dpf. The nearest gene is indicated below each track. (E) Violin plots showing cell type-enriched expression of genes proximal to corresponding CREs. *gata3* was enriched in the 3 dpf scRNA-seq dataset but exhibited sparse expression, so the 2 dpf dataset is shown for visualization. All remaining genes are shown using 3 dpf scRNA-seq data (Raj et al. 2020). (F) In vivo enhancer activity of selected CREs in 3 dpf larvae. Maximum-intensity projections of confocal images are shown. Ratios in the lower left corner indicate the number of eGFP+ larvae relative to total mCherry+ transgenic larvae. Regions of interest are outlined with dashed white lines. White arrowheads mark the hindbrain (CRE72) and RGC axons at the optic chiasm (CRE77). The bottom CRE77 panel shows a higher-magnification view of the region above. (G) Colocalization of CRE77 reporter activity (left) with the Tg(*isl2b-gata2aBP*:mKate) line (right). Bottom panel, merged confocal image. White arrows denote representative cells with overlapping signal in yellow. White arrowhead represents RGC axonal tracts crossing into the brain. Scale bars, 100 μm.

Putative intergenic CREs were cloned upstream of a minimal promoter (Kemmler et al. 2023) to test their ability to drive eGFP expression in vivo (Fig. 4B, C). We classified intergenic CREs as proximal (1-10 kb from the transcription start site, TSS) and distal (> 10 kb from the TSS). Proximal elements were subsequently prioritized because they more frequently produced robust reporter activity in initial screens. Out of 23 proximal CREs tested, 11 reproducibly drove eGFP expression in spatial patterns that mirror nearby marker genes, whereas only one of the seven distal CREs showed detectable enhancer activity (Supplemental Table S9, Fig. 4D-F, Supplemental Fig. S9). For example, CRE58 drove reporter expression in the optic tectum, consistent with its accessibility near the tectally expressed *gata3*. Two enhancers, CRE63 and CRE64, were active in radial glia, matching the expression of the nearby radial glia marker *slc1a3b*. Interestingly, although CRE72 was accessible across multiple cell types, our pipeline identified it as preferentially enriched in hindbrain populations. Reporter assays confirmed enhancer activity in the hindbrain, consistent with expression of the nearest gene, *meis2b*, in hindbrain cells. CRE77, located near *isl2b*, drove reporter expression in RGCs, where *isl2b* is also expressed, and labeled their axonal projections through the optic chiasm. To confirm RGC-specific activity, the CRE77 plasmid was injected into embryos from Tg(*isl2b-gata2aBP*:mKate) line, which pairs a 7.6 kb *isl2b* fragment (Fredj et al. 2010) with a gata2a basal promoter and marks RGCs. Colocalization of eGFP and mKate expression validated CRE77 activity within the RGC population (Fig. 4G). We further confirmed retinal eGFP expression in CRE77 F1 transgenic animals (Supplemental Fig. S10). Although larger regulatory fragments from all these genes have been used in transgenic reporters (Fujimoto et al. 2011; Callahan et al. 2019; Guerra et al. 2018; Chen et al. 2020; Gall et al. 2025; Fredj et al. 2010; Pittman et al. 2008), the specific cis-regulatory modules driving these expression patterns have not been characterized. Together, these validations demonstrate the power of our resource for systematic CRE discovery and functional annotation in the nervous system.

### Dissection of *slc1a3b* enhancer regulatory architecture

To determine whether this resource could be used to resolve enhancer architecture at high resolution, we focused on the *slc1a3b* locus (GLAST/SLC1A3 ortholog), encoding a glutamate transporter expressed in radial glia and astroglia. Prior studies used larger regulatory fragments (9.5 kb and 3 kb) to label *slc1a3b*-expressing cells (Chen et al., 2020; Gall et al., 2025), but the discrete regulatory elements underlying this activity were not defined. CRE63 is contained within both fragments, whereas CRE64 partially overlaps the 9.5 kb fragment but is absent from the 3 kb construct (Fig. 5A). Both elements are marked by H3K27ac and H3K4me1 (Baranasic et al. 2022), consistent with active enhancer chromatin. In addition, CRE63 and CRE64 coincide with conserved non-coding elements across teleost species, suggesting evolutionary constraint and functional importance (Supplemental Fig. S11).

**Figure 5.**
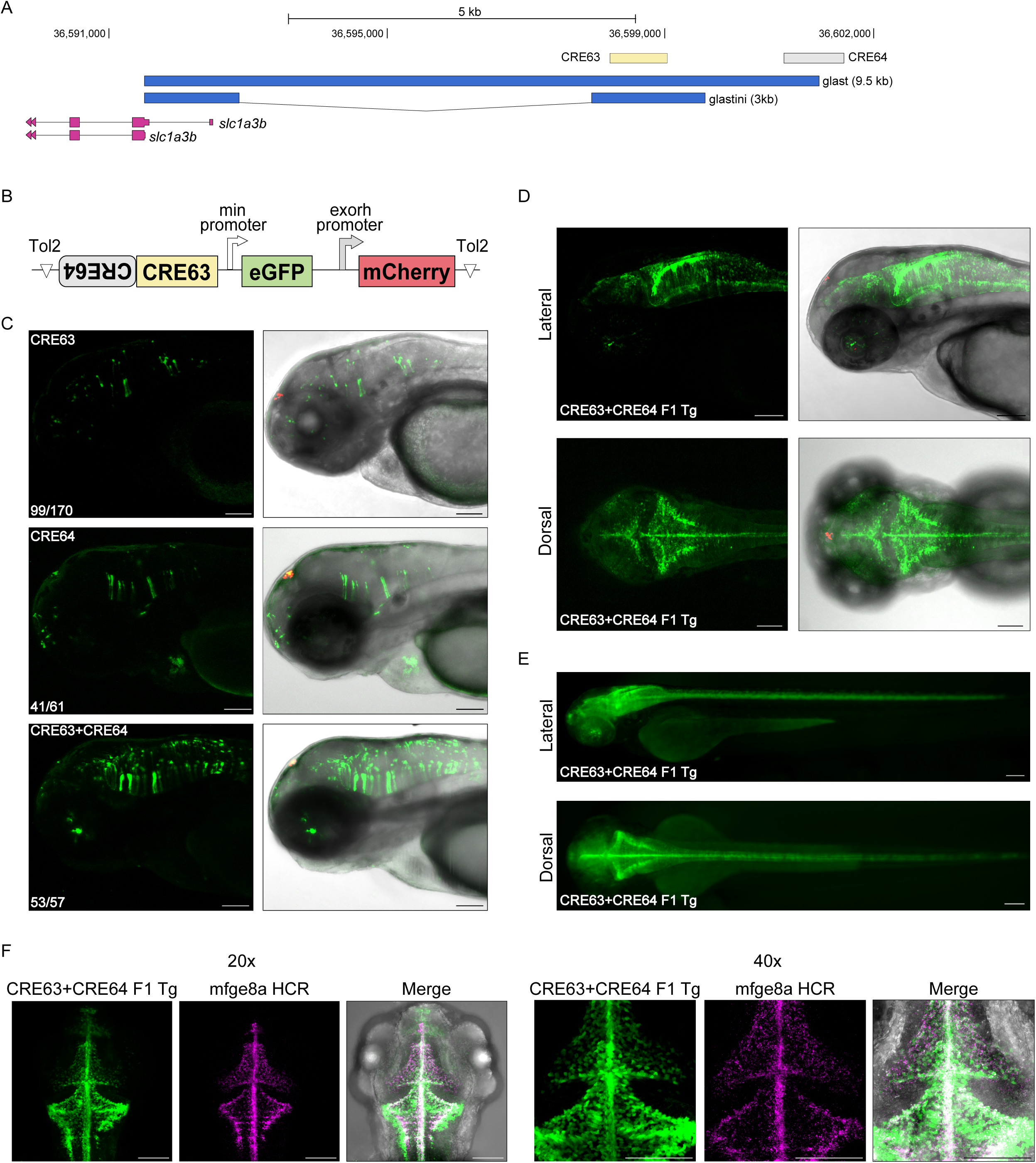
Combinatorial activity of CRE63 and CRE64 drives radial glial reporter expression. (A) Genomic view of the *slc1a3b* locus. Previously published promoter fragments (9.5 kb and 3 kb) and two candidate enhancer elements identified in this study, CRE63 and CRE64, are shown. Scale bar, 5 kb. (B) Schematic of the enhancer reporter construct used to test combinatorial activity. CRE63 and CRE64 were cloned individually or concatenated upstream of a minimal promoter driving eGFP. (C) Representative confocal images of 3 dpf zebrafish expressing CRE63, CRE64, or CRE63+CRE64 reporter constructs. Maximum-intensity projections are shown. Ratios in the lower left corner indicate the number of eGFP+ larvae relative to total mCherry+ transgenic larvae. (D) Representative confocal images of 3 dpf CRE63+CRE64 F1 transgenic zebrafish. (E) Whole-body fluorescence images of 3 dpf CRE63+CRE64 F1 transgenic fish. (F) Confocal images of 3 dpf CRE63+CRE64 F1 transgenic fish showing eGFP expression (green) and *mfge8a* transcripts detected by HCR (magenta). Merged images show overlap of both signals, with representative double-positive cells appearing white. Left panels, 20x objective. Right panels, 40x objective. Scale bars in C-F, 100 μm.

To test whether CRE63 and CRE64 functionally interact, we concatenated the two elements in our reporter construct (Fig. 5B). When tested individually, each element was sufficient to drive reporter activity in a radial-glia associated pattern (Fig. 5C). Strikingly, concatenation of CRE63 and CRE64 resulted in a marked increase in reporter activity exceeding that of either element alone (Fig. 5C, Supplemental Fig. S12A-C), consistent with combinatorial enhancer activity. CRE63+CRE64 transgenic F1 animals showed robust eGFP expression in a radial glia-associated pattern along the brain midline and diamond-shaped hindbrain ventricle, and extending down the spinal cord (Fig. 5D, 5E), closely mirroring the previously established *slc1a3b* transgenic line derived from the 9.5 kb fragment (Supplemental Fig. S12D). We also noted weak labeling in the retina of F1 animals, potentially corresponding to Müller Glia cells. To confirm expression in radial glia cells, CRE63+CRE64 F1s were probed by in situ hybridization chain reaction to detect mRNA expression of the radial glia marker *mfge8a*. Extensive overlap of eGFP and *mfge8a* signals in the midbrain and hindbrain confirmed enhancer activity in radial glia cells (Fig. 5F).

Next, we sought to identify important regulatory sequences within the enhancer fragments. We first performed deletion analysis on CRE63 in ~200 bp increments (Fig. 6A). Deletion of the first 200 bp (Δ200) retained reporter activity. However, larger deletions (Δ400 and Δ600) abolished expression, localizing essential sequences to the deleted region between Δ200 and Δ400 (min200-400). Injection of the min200-400 fragment alone did not restore eGFP expression, indicating it is required but not sufficient for full enhancer function. Motif scanning of this interval with FIMO identified multiple candidate TF binding sites, including predicted binding sites for TEAD1 and STAT3, which are associated with Hippo/YAP and JAK/STAT signaling, respectively (Fig. 6A). An additional predicted STAT3 binding site is located further downstream in CRE63. Tead1 expression is restricted to intermediate progenitors and radial glia cells in the developing mouse brain (Pelenyi et al. 2024). Stat3 has an established role in radial glia proliferation and injury-induced activation in the zebrafish central nervous system (Shimizu et al. 2021; Shimizu and Kawasaki 2021). Motif analysis of CRE64 identified predicted binding sites for SOX9, ETV5, POU3F3, and STAT1::2. Notably, these predicted sites overlap a region of high conservation across teleost species (Supplemental Fig. S13). ChromVAR analysis showed strong enrichment of tead1b, stat3, sox9a, pou3f3b, and stat1b+stat2 motif accessibility in radial glia and weak enrichment for etv5b (Supplemental Fig. S14A). Consistent with a functional role in radial glia, scRNA-seq data confirmed expression of *tead1b*, *stat3*, *sox9a*, *etv5b*, and *pou3f3b* in the radial glia cluster at 3 dpf (Fig. 6B). However, we did not detect *stat1b* or *stat2* expression in this cluster. Together, these findings identify candidate transcriptional regulators of CRE63 and CRE64 activity (Fig. 6C).

**Figure 6.**
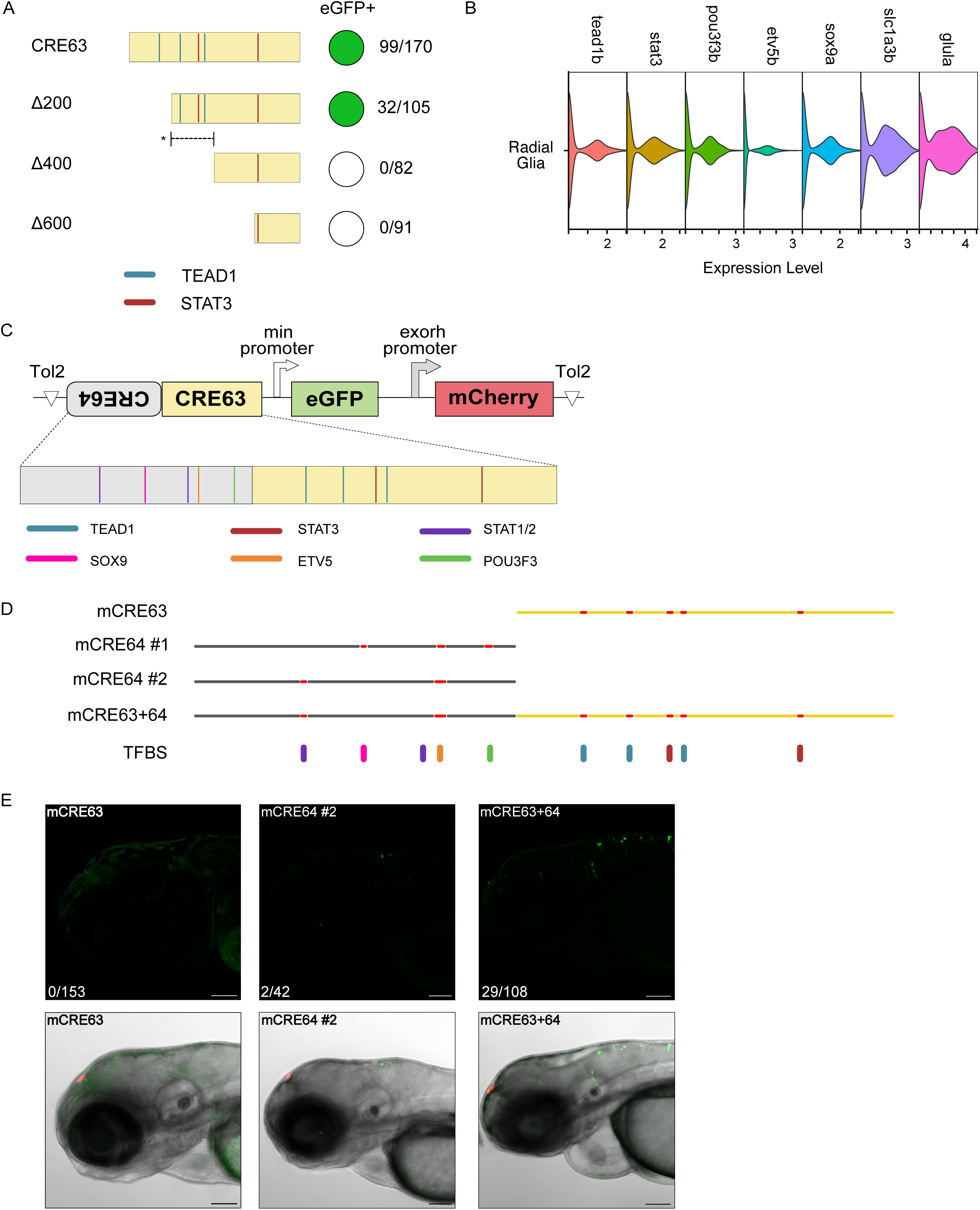
Dissection of modular enhancer architecture at the *slc1a3b* locus. (A) Schematic of CRE63 deletion constructs. Ratios to the right indicate the number of eGFP+ larvae relative to total mCherry+ transgenic larvae. Predicted binding sites for TEAD1 and STAT3 are indicated. Asterisk represents a critical regulatory region between Δ200 and Δ400 (min200-400) required for enhancer activity. (B) Violin plots of candidate TFs associated with CRE63 and CRE64 and canonical radial glia markers (*slc1a3b*, *glula*) from 3 dpf scRNA-seq data. (C) Schematic of combinatorial CRE63+CRE64 enhancer reporter construct with predicted TF binding sites shown. (D) Schematic of CRE63 and CRE64 motif perturbations. Yellow and grey lines indicate CRE63 and CRE64 sequences, respectively. Red segments indicate perturbed motifs in each mutant construct. TFBS, predicted transcription factor binding sites. (E) Representative confocal images of 3 dpf zebrafish expressing mutated CRE reporter constructs. Maximum-intensity projections are shown. Ratios in the lower left corner indicate the number of eGFP+ larvae relative to total mCherry+ transgenic larvae. Scale bars, 100 μm.

### Motif perturbation

To test the functional importance of the identified candidate binding sites, we performed motif perturbation experiments. We generated reporter constructs harboring mutations in different combinations of CRE63 and CRE64 motif sequences by randomly scrambling the nucleotides within each motif (Fig. 6D, Supplemental Fig. S14B). This approach largely preserved overall enhancer length and nucleotide composition while disrupting the native motif sequences. Within CRE63, mutation of all identified STAT3 and TEAD1 motifs abolished eGFP expression (Fig. 6E). Within CRE64, mutation of predicted SOX9, POU3F3, and ETV5 motifs reduced but did not abolish eGFP reporter expression (Supplemental Fig. S14C). We suspected that the initial ETV5 mutation did not fully disrupt ETS-family binding potential. Therefore, we generated a second CRE64 mutant reporter in which the ETV5 mutated region was extended and combined with mutation of one of the two STAT1::2 sites. This construct abolished eGFP expression (Fig. 6E). Finally, we combined the CRE63 and CRE64 mutations in a concatenated enhancer. This mutant reporter showed only sparse labeling of a few cells compared with the strong and extensive activity of the wildtype concatenated sequence. Together, these results demonstrate that CRE63 and CRE64 enhancer activity depends on multiple cooperative cis-regulatory inputs.

### Shared cis-regulatory inputs in human *SLC1A3*

Finally, we asked whether the candidate *slc1a3b* regulatory factors identified in zebrafish are also implicated in human *SLC1A3* regulation. To address this, we leveraged publicly available ENCODE enhancer annotations and published ChIP-Seq datasets to interrogate ~10 kb upstream of the human *SLC1A3* locus (Supplemental Fig. S15). Within this region, we identified multiple annotated enhancers containing STAT3, TEAD1, ETV5, and STAT1::2 binding motifs. Notably, a predicted STAT3 motif within an annotated enhancer overlaps STAT3 ChIP-seq peaks, supporting direct binding activity. In addition, a STAT3 motif near the promoter also coincides with STAT3 ChIP-seq signal, and the ETV5 motif resides within a highly conserved vertebrate sequence. Together, these orthogonal lines of evidence support the possibility that similar TF regulatory inputs contribute to human SLC1A3 control.

## DISCUSSION

We present a single-cell chromatin accessibility dataset of the zebrafish brain and retina spanning larval, juvenile, and adult stages. While regulatory landscapes during early zebrafish embryogenesis have been extensively profiled and resources such as DANIO-CODE (Baranasic et al. 2022) and Zebrahub (Kim et al. 2024; Lange et al. 2024) have substantially expanded annotation of the zebrafish regulatory landscape, chromatin architectures and cis-regulatory logic during post-embryonic neural expansion remain largely unexplored. To our knowledge, only one zebrafish brain-specific scATAC-seq dataset has been reported to date, identifying 25 chromatin-defined clusters in the adult brain (Yang et al. 2020). By resolving >200 cis-regulatory states across neural cell types and post-embryonic timepoints, this resource substantially expands the resolution of the zebrafish neural chromatin landscape. It enables cell type-, region-, and stage-resolved analysis of chromatin remodeling, inference of TF regulatory programs, and identification of candidate enhancers across post-embryonic development.

Our analyses reveal two overarching principles governing vertebrate neural gene regulation. First, neural cell types and brain regions undergo substantial remodeling of their cis-regulatory landscapes across the profiled post-embryonic periods. Stage-enriched DA peaks reflected both the gain of later-stage accessible elements and the loss of elements that were preferentially accessible earlier in development. The functional associations of these regions were consistent with this developmental progression: larval-enriched DA peaks were linked to programs, including cell adhesion, axon guidance, and Notch signaling; whereas, adult-enriched DA peaks were associated with mature neuronal functions, including synaptic organization, circadian regulation, and metabolic homeostasis. Second, our datasets provide a framework for defining cell type- and region-specific regulatory programs. Integration of transcriptomic and chromatin accessibility data linked TF expression and gene activity with enrichment of cognate TF motifs, identifying candidate regulators associated with distinct neural chromatin states. Comparisons across timepoints further revealed that the temporal stability of these regulatory programs varies across neural populations. Some cell types and regions retained relatively stable motif signatures, whereas others underwent substantial divergence, indicating that post-embryonic maturation involves both preservation of core regulatory identities and stage-dependent reconfiguration of TF-associated regulatory programs.

Furthermore, our datasets enable systematic discovery and validation of candidate enhancers across the zebrafish brain and retina. Chromatin accessibility is a hallmark of active CREs and has been used successfully to identify functional enhancers across zebrafish tissues (Quillien et al. 2017; Thompson et al. 2020). Consistent with this, we validated enhancer activity for 11 of 23 tested proximal CREs, corresponding to a success rate of nearly 50%, indicating that screening only a small number of prioritized candidates is often sufficient to identify a functional element. This success rate may be further improved by incorporating orthogonal epigenetic information when available. Importantly, reporter context may also influence apparent CRE activity as enhancer output can depend on the minimal promoter used (Quillien et al. 2017; Kemmler et al. 2023). Therefore, lack of eGFP expression in our assay does not necessarily indicate that a candidate CRE is non-functional. We generally linked accessible regions to the nearest annotated gene and used scRNA-seq expression as supporting evidence for putative target assignment. However, enhancer-promoter interactions can span considerable genomic distances and do not always involve the nearest gene. Finally, we focused on validating intergenic accessible regions, which represents only a subset of the regulatory landscape. In the future, our approach can be extended to intronic and other classes of CREs.

These findings highlight the utility of our single-cell chromatin accessibility resource for enhancer discovery in the zebrafish nervous system. Traditional zebrafish enhancer discovery has relied on BAC transgenesis, promoter cloning, or enhancer trapping. While powerful, these approaches often lack resolution and mechanistic specificity. BAC reporters can span tens to hundreds of kilobases and recapitulate expression without isolating the discrete enhancers responsible. For example, a *meis2b* BAC transgenic line incorporated 139 kb of upstream genomic sequence (Guerra et al. 2018), whereas our data identifies a 978 bp accessible fragment sufficient to label hindbrain cells. Similarly, 14.7 kb and 7.6 kb fragments were used to generate RGC reporters (Pittman et al. 2008; Fredj et al. 2010), whereas we identified a 723 bp CRE that labels these cells. Enhancer trapping provides an unbiased route to expression drivers but is stochastic, labor-intensive and requires secondary mapping to define the regulatory element and its target gene (Trinh and Fraser 2013). In contrast, our approach yields compact, gene-linked regulatory candidates across cell types and regions at genome-wide scale, accelerating functional annotation of the zebrafish regulatory genome. This will be valuable for generating transgenic lines that mark specific neural populations, reducing the need to clone large regulatory fragments.

Finally, using the *slc1a3b* locus we demonstrate how this resource can be leveraged to dissect enhancer architecture in vivo. We identified two discrete regulatory elements that act combinatorially to drive *slc1a3b* expression in radial glia cells and defined candidate TF binding motifs required for activity of each enhancer individually and in combination. These findings implicate JAK/STAT-, Hippo/YAP-, and ETS-family-associated regulatory inputs in controlling radial glia-specific *slc1a3b* expression. Although *slc1a3b* is not detected in zebrafish Müller glia, our enhancer constructs showed weak retinal activity, suggesting that additional repressive elements may contribute to restricting endogenous *slc1a3b* expression. SLC1A3 (GLAST) is expressed in mammalian radial glia and astroglial populations, including Müller glia, and plays key roles in glutamate homeostasis and neural function. Strikingly, the upstream regulatory landscape of human *SLC1A3* contains predicted TF binding sites that parallel the candidate regulatory inputs identified at the zebrafish *slc1a3b* locus. Consistent with this, modulation of JAK/STAT signaling alters *SLC1A3* expression in astrocytes (Wang et al. 2016; Feng et al. 2015), while ETV5 overexpression in mammalian radial progenitors increases the proportion of Slc1a3-positive astrocyte precursors (Li et al. 2012). Although direct binding by these TFs at human enhancers remains to be tested, these observations raise the possibility that components of the regulatory network controlling *slc1a3b/SLC1A3* are conserved across vertebrates.

In summary, this work illustrates how single-cell chromatin datasets can move beyond descriptive profiling to enable discovery, functional validation, and cross-species comparison of cis-regulatory elements. Together, our study establishes a foundational resource for defining enhancer logic across post-embryonic neural development with a direct impact on identification of candidate regulatory elements and factors for targeted perturbation in the vertebrate nervous system.

## MATERIAL AND METHODS

### Zebrafish husbandry

All zebrafish work was performed under protocol numbers 807110 and 807259, which were approved by the University of Pennsylvania’s Office of Animal Welfare and Institutional Animal Care and Use Committee (IACUC).

### scATAC-seq

Details of single-nuclei isolation steps are provided in Supplemental Methods. Samples were run using the 10x Chromium platform according to the manufacturer’s instructions (Chromium Next GEM Single Cell ATAC Library & Gel Bead Kit v2), targeting ~6000 nuclei per channel. Pooled libraries were sequenced using either Illumina NovaSeq 6000 or Illumina NovaSeq X Plus.

To supplement cell numbers and provide a benchmark for data quality, we incorporated public scATAC-seq data from (Yang et al. 2020). Raw sequencing reads for adult zebrafish brain samples (GSM4662087 and GSM4662088) were retrieved from the NCBI Gene Expression Omnibus (GEO) under accession GSE134055. Raw sequencing data were processed using the Cell Ranger ATAC pipeline (v2.1.0, 10x Genomics). FASTQ files were aligned with the cellranger-atac count command to a reference genome built from the zebrafish GRCz11 (danRer11) primary assembly and UCSC formatted Ensembl gene annotations (danRer11.ensGene.gtf). See Supplemental Methods for details on filtering steps and cell type annotation.

### Creation of a unified consensus peak set

Peak sets originally identified by the CellRanger ATAC pipeline for each stage were imported as GenomicRanges objects and merged into a non-redundant master set. To remove potential artifacts, we applied genomic filters to the combined peaks (Minimum width: 20 bp, Maximum width: 10,000 bp). The consensus peak set was then used to re-quantify chromatin accessibility across all stages. For each dataset, a cell-by-peak feature matrix was generated using the FeatureMatrix function. Chromatin assays were annotated using the Ensembl Release 112 GTF (Danio_rerio.GRCz11.112.chr.gtf). A ChromatinAssay was then created for each sample and incorporated into a Seurat object for downstream analysis.

### Identification of stage-specific differentially accessible regions

To assess changes in chromatin accessibility across post-embryonic stages, scATAC-seq datasets from all timepoints were merged using Seurat. Non-neuronal clusters were excluded. Following normalization, the merged object was batch corrected using RunHarmony (Harmony package). Differential accessibility was assessed through pairwise comparisons between each stage using FindMarkers (Seurat). Peaks were considered differentially accessible if the average log2 fold-change >1.0 and the Benjamini–Hochberg adjusted p-value <0.05. For significant peaks, mean fragment counts per timepoint were computed using AggregateExpression, log-transformed, and z-score normalized for visualization.

### Cell type-resolved changes in chromatin accessibility

To identify temporal changes in peak accessibility, retinal cells were subset, integrated, and annotated as described above. A population of 21 cells lacking clear cluster assignment and canonical marker expression was excluded prior to analyses. The integrated retinal object was partitioned by cluster identity, and pairwise timepoint comparisons were performed using FindMarkers (Wilcoxon rank-sum test). Comparisons were restricted to groups containing at least 10 cells. Peaks detected in fewer than 10% of cells in either group (min.pct = 0.1) were excluded, and an initial log2 fold-change threshold of 0.5 was applied to improve computational efficiency. Final significance thresholds were set at average log2 fold-change >1.0 and adjusted p-value <0.05. For peaks identified as differentially accessible in multiple timepoint comparisons within a cluster, only the instance with the highest log2 fold-change was retained to generate a non-redundant peak list per cluster. The same analytical framework was applied to brain populations. To assess confounding effects of cluster size on DA peak detection, Spearman correlation and linear regression analyses were performed between cell number per cluster and DA peak counts.

### Peak-gene linkage analysis

Peak-gene links were computed on the integrated retina and brain scATAC objects using the Signac and Seurat packages in R. Prior to linkage analysis, the gene activity assay was log-normalized using NormalizeData, and highly variable features were identified with FindVariableFeatures followed by ScaleData. Chromatin accessibility profiles were analyzed using the ATAC assay with the *Danio rerio* danRer11 reference genome (BSgenome.Drerio.UCSC.danRer11). GC-content and other sequence-level statistics required for background correction were computed for all peaks genome-wide using RegionStats. Peak-gene links were then inferred with LinkPeaks, pairing each ATAC peak with genes whose accessibility co-varied across single nuclei within a maximum genomic window of 500 kb from the TSS. Only peaks detected in a minimum of 10 cells were considered. Pearson correlation coefficients and associated empirical *p*-values were computed against a background model of matched GC-content and peak accessibility. Peak-gene associations were retained if they met a correlation coefficient threshold of |score| ≥ 0.05 and an empirical p-value < 0.05.

### Identification of genes linked to differentially accessible regions

Differentially accessible regions (DARs) identified as enriched at 3 dpf or adult stages in the cell-type resolved analysis were intersected with the peak-gene link table described above, yielding a set of stage-specific, cluster-resolved peak-gene pairs. Peaks lacking an assigned gene link were excluded from gene ontology analyses.

### Gene Ontology enrichment analysis

Gene Ontology (GO) enrichment analysis was performed using the clusterProfiler package (v4.18.4) with the *Danio rerio* annotation database (org.Dr.eg.db). A background gene universe was constructed from gene annotations derived from the HOMER-annotated peak set, retaining only genes associated with peaks accessible in at least 10% of nuclei in either the retina or neuron dataset. These gene symbols were mapped to Entrez IDs via bitr (fromType = “SYMBOL”, toType = “ENTREZID”). Gene symbols associated with DAR-peak links were similarly converted to Entrez IDs and filtered to retain only genes present in the predefined universe. GO enrichment of Biological Process (BP) terms was performed independently for each cluster–stage group using enrichGO with the following parameters: ontology = “BP”, *p*-value adjustment method = Benjamini–Hochberg (BH), *p*-value cutoff = 0.1, and *q*-value cutoff = 0.20. For visualization, the top three GO terms per cluster–stage combination were selected based on adjusted *p*-value, and enrichment scores were expressed as −log10(FDR-adjusted *p*-value).

### Motif enrichment analysis

Motif annotations were added to Signac objects from each timepoint using AddMotifs with TF position weight matrices from the DANIO-CODE consortium (Baranasic et al. 2022). Motif accessibility deviations were computed using RunChromVAR, generating a motif assay containing per-cell deviation scores. Differentially active motifs across clusters were identified with FindAllMarkers. Motifs were ranked by average difference per cluster, and mean deviation scores were calculated using AverageExpression. Top differentially active motifs were z-score normalized.

### Co-embedding of scRNA-seq and scATAC-seq data

Gene expression was imputed onto scATAC-seq cells from 3 dpf using previously identified integration anchors. Expression values for the top 5,000 variable genes from the 3 dpf scRNA-seq reference were transferred to the scATAC-seq query. The scRNA-seq and imputed scATAC-seq datasets were merged, and principal component analysis was performed on shared variable features. Co-embedded objects were plotted on the same UMAP, enabling joint visualization of gene expression, chromatin accessibility, and motif enrichment profiles.

### Jaccard similarity analysis

Motifs were identified using FindMarkers and considered significant if they exhibited an adjusted p-value <0.05 and an average difference exceeding the median threshold (0.8). For each cell lineage at each timepoint, a binary matrix of significant motifs was generated. Pairwise Jaccard similarity scores were computed using the proxy package in R and visualized as heatmaps ordered by lineage and developmental stage.

### Identification of putative cis-regulatory elements

DARs from 3 dpf scATAC-seq data were annotated using HOMER’s (Heinz et al. 2010) *annotatePeaks* function to assign proximal genes. DARs were retained if their associated gene was also identified as an upregulated differentially expressed gene (DEG) in 3 dpf scRNA-seq data. The resulting set was filtered to include only intergenic regions detected in at least 10% of cells within the cluster and exhibiting a fold change > 1.5. Remaining DARs were ranked by percent difference and fold change. Putative CREs were classified based on distance from the transcription start site (TSS): regions located 1–10 kb upstream of the TSS were designated proximal, whereas those located more than 10 kb upstream were classified as distal.

### Enhancer reporter assays

Reporter constructs containing putative CREs were generated using a modification of vector pCK083 desmaMCS:minpromEGFP, exorh:mCherry (gift from Christian Mosimann; Addgene plasmid #200016). The *desma* element was removed using a Q5 site-directed mutagenesis kit, resulting in plasmid p228. Candidate CREs were PCR amplified from genomic DNA and inserted into p228 by Gibson assembly following AgeI digestion. Primer sequences are reported in Supplemental Table S9. Reporter plasmids were co-injected with Tol2 mRNA into one-cell-stage embryos, as described previously (Raj et al. 2018). Embryos were treated with 1-phenyl-2-thiourea (PTU) before 24 hpf and screened for eGFP expression at 72 hpf. See Supplemental Methods for further details.

### Imaging

Fluorescence imaging of 3 dpf larvae was performed using a Leica M165 FC. Confocal imaging was performed using a 10x lens on a Zeiss LSM 880 laser scanning confocal microscope. Images were processed in Fiji (ImageJ version 1.54p, Java 1.8.0_172). HCR RNA-FISH was performed as described previously (Siniscalco et al. 2024). Probes are provided in Supplemental Table S10.

### Motif analysis

Motif scanning of *slc1a3b* and *SLC1A3* loci were performed using FIMO (MEME Suite v5.5.9) with vertebrate motifs from the JASPAR 2026 database. TFs were prioritized as candidate regulators if motifs met q-value < 0.05 and the corresponding genes were expressed in radial glia in the 3 dpf scRNA-seq dataset.

### Motif Perturbations

Candidate TF binding sites within CRE63 and CRE64 were disrupted by randomly shuffling motif sequences. For CRE63, nucleotides within the predicted STAT3 and TEAD1 binding motifs were altered. For CRE64, motif perturbation was performed in two iterations. In the first iteration, predicted POU3F3, ETV5, and SOX9 binding sites were disrupted. In the second iteration, ETV5 motif and a predicted STAT1::STAT2 motif were disrupted. All mutant enhancer sequences were synthesized as gBlocks and cloned into the p228 reporter backbone using the same strategy as wildtype enhancer constructs.

### Quantification of mosaic F0 eGFP-positive fish

The ratio of eGFP to mCherry transgene-positive larvae were scored. eGFP-positive fish were defined by clear GFP expression across numerous cells within the expected region. For motif perturbation experiments, fish were considered eGFP-positive if more than two brightly labeled eGFP-expressing cells were detected within the expected region and clear radial glia-like morphology could be seen.

## DATA ACCESS

All raw and processed sequencing data generated in this study have been submitted to the NCBI GEO, under accession number GSE324435. R scripts used for data processing are publicly available on GitHub at: https://github.com/BushraRaj-Lab/Greenslade_2026.

## COMPETING INTEREST STATEMENT

The authors declare no competing interests.

## ACKNOWLEDGEMENTS

We thank the UPenn zebrafish facility staff for assistance with animal husbandry and the UPenn Cell and Developmental Biology Microscopy Core (RRID SCR_022373). We thank Maggie Zhang for experimental assistance, Roshan Perera for bioinformatic assistance, and Philip Campbell (UPenn) for sharing embryos from the (−9.5kb)slc1a3b:myrGCaMP6s-P2A-H2AmCherry transgenic line. The Tg(isl2b-gata2aBP:mKate) line was a gift from Michael Granato (UPenn) and requests for this line should be directed to the Granato lab. This work was supported by a National Science Foundation GRFP grant DGE-2236662 to J.E.G., the National Institutes of Health grants R00HD098298 and DP2NS131787 to B.R., and startup funds from the University of Pennsylvania to B.R. A total of $480,000, or 80 percent of this project was financed with federal funds. A total of $120,000, or 20 percent of the project was financed with non-governmental funds. The content is solely the responsibility of the authors and does not necessarily represent the official views of the National Institutes of Health.

## Author contributions

B.R. and J.E.G designed the study. B.R. and J.E.G. wrote the manuscript with input from all authors. J.E.G. collected and analyzed scATAC-seq data and generated figures. H.V. and J.E.G performed confocal microscopy. J.E.G and M.L.R. cloned constructs and performed microinjections.

## Notes

### Competing Interest Statement

The authors have declared no competing interest.

### Summary of Updates

1. We added new biological insights related to Fig. 2 and Fig. 3. 2. We added F1 characterization of two enhancers. 3. We added cell-type validation to support enhancer activity in retinal ganglion cells and brain radial glia. 4. We mutated predicted transcription factor motifs in the CRE63+64 enhancer reporter construct. 5. We uploaded our data to the interactive UCSC Cell Browser website. 6. We updated Supplemental files.

