## Supplemental Material for "Single-cell chromatin profiling reveals dynamic regulatory logic and enhancer elements in brain and retina development"

### Supplemental Methods

#### Single-nuclei dissociation

Wildtype zebrafish (TL/AB) at 3 dpf, 21 dpf and adult (5 months post-fertilization, mpf) stages were used for scATAC-seq. For 3 dpf samples, the head region including the eyes were dissected, and heads from ~10 larvae were pooled per collection. For 21 dpf and adult stages, brains and eyes were dissected separately and pooled (21 dpf: 3 brains and 4 eyes; adult: 1 brain and 2 eyes). All dissections were performed in cold Neurobasal media supplemented with B27 and tissues were pooled in 1 mL of cold Neurobasal/B27 solution. Pooled 3 dpf heads in 1 mL Neurobasal/B27 were immediately transferred to a pre-chilled Dounce homogenizer and homogenized by four strokes with pestle A followed by four strokes with pestle B, until no visible tissue fragments remained. The homogenate was transferred to a 2.0 mL tube and centrifuged at 500 × g for 4 min at 4°C. The pellet was resuspended in 100 µL of 0.1× lysis buffer (10 mM Tris-HCl pH 7.4, 10 mM NaCl, 3 mM $MgCl_2$, 1% BSA, 0.1% Tween-20, 0.1% Nonidet P40, 0.01% digitonin) by pipetting ten times, then incubated on ice for 5 min. Following lysis, 1 mL of wash buffer (10 mM Tris-HCl pH 7.4, 10 mM NaCl, 3 mM $MgCl_2$, 1% BSA, 0.1% Tween-20) was added and mixed by pipetting five times. The nuclei suspension was sequentially filtered through 35 µm and 20 µm strainers, then centrifuged at 500 × g for 1 min at 4°C. Nuclei were resuspended in 1× nuclei buffer (10x Genomics), counted using a hemocytometer, and diluted to ~2,500 nuclei/µL for downstream processing. Nuclei isolation for 21 dpf samples was performed similarly, except tissues were homogenized with approximately eight strokes each using pestles A and B. In addition, post-wash filtration was performed sequentially through 20 µm and 10 µm strainers.

For adult zebrafish samples, an adjusted nuclei isolation protocol was used to reduce cellular debris in the final suspension. The lens and cornea were removed from adult eyes prior to processing. Brain and eye tissues were resuspended in 1 mL of 0.1× lysis buffer and transferred to a pre-chilled Dounce homogenizer. Tissues were homogenized with ~12 strokes using pestle A followed by 12 strokes using pestle B. The homogenate was incubated on ice for 5 min. The suspension was filtered through a 35 µm strainer, transferred to a 2.0 mL tube, and centrifuged at 500 × g for 4 min at 4°C. The pellet was resuspended in 500 µL wash buffer and mixed with 900 µL of 1.8 M sucrose solution (15.4 g sucrose dissolved in 25 mL wash buffer). An additional 500 µL of sucrose solution was carefully layered on top to form a cushion. Samples were centrifuged at 13,000 × g for 5 min at 4°C to pellet nuclei. The nuclei pellet was resuspended in 500 µL wash buffer, then centrifuged again at 500 × g for 2 min at 4°C. Final nuclei were resuspended in 1× nuclei buffer, counted using a hemocytometer, and diluted to ~ 2,500 nuclei/µL for downstream scATAC-seq processing.

For the 3 dpf timepoint, samples were collected on two separate days, with two technical replicates per day, yielding four total libraries. For the 21 dpf timepoint, samples were collected across three days, generating four brain libraries and two eye libraries. Adult samples were collected on a single day, with two technical replicates each for both eye and brain samples.

**scATAC-seq quality control and filtering**

Libraries from the same developmental timepoint were aggregated using cellranger-atac aggr. The resulting peak-barcode matrices and fragments files were used for downstream analysis. Standard quality control procedures implemented in the Signac package were used to remove low-quality nuclei from each dataset. Nuclei were retained if they met the following criteria: >2,000 fragments in peaks, >60% of reads mapping to peaks, nucleosome signal <4, and transcription start site (TSS) enrichment score >2.

**Cell type annotation**

To annotate cell types, we integrated the 3 dpf scATAC-seq dataset with previously published whole-head 3 dpf scRNA-seq data (Raj et al. 2020). Brain cells from the 21 dpf and adult scATAC-seq datasets were integrated with a published 21 dpf brain scRNA-seq reference (Siniscalco et al. 2024). Retinal and other eye cells from the 21 dpf and adult scATAC-seq datasets were separately integrated with a published 15 dpf scRNA-seq dataset containing eye tissue (Raj et al. 2020). Integration was performed using FindTransferAnchors and cell type labels were projected from the scRNA-seq reference onto scATAC-seq cells using TransferData. Cluster annotations were assigned based on the strength of the predicted label scores and the proportion of cells within each cluster sharing the same transferred identity. Final annotations were refined through iterative subclustering and manual inspection of gene activity scores for canonical marker genes.

**Concatenation of CRE63 and CRE64 elements**

CRE63 and CRE64 were combined into a single tandem construct via PCR. Individual CREs were first PCR amplified from existing plasmid templates using the following primers: CRE63fwd: tgagggtgagtaaatggtgag, CRE63rev: ggggctaatgattctgacatg, CRE64fwd: ggacatcttgcactcttgtc, CRE64rev: atgtcaaacccttgacgtcc. A second PCR was performed using primers that introduced a complementary fusion bridge between the two elements: CRE63fwd-2: ggacgtcaagggtttgacaTGAGGGTGAGTAAATGGTGAG, CRE63rev-2: ggggctaatgattctgacatg,

CRE64fwd-2: ggacatcttgcactcttgtc, CRE64rev-2: ctcaccatttactcaccctca-TGTCAAACCCTTGACGTCC, where lowercase letters indicate the bridging overhangs. The CRE fragments were then fused in a two-stage overlap extension-PCR. Fragments were first annealed with 12 cycles without any primers and then amplified with 30 cycles using the primers Cre63rev-3: gaattcgcccttcccggggtcgacaggggctaatgattctgacatg and Cre64fwd-3: ggctccgaattcgcccttgatatcaggacatcttgcactcttgtc.

Motif-perturbed mCRE63+CRE64 constructs were generated using the same overlap extension PCR strategy, except that synthesized gBlocks (Integrated DNA Technologies) containing the mutated enhancer sequences were used as templates for the initial PCR amplifications.

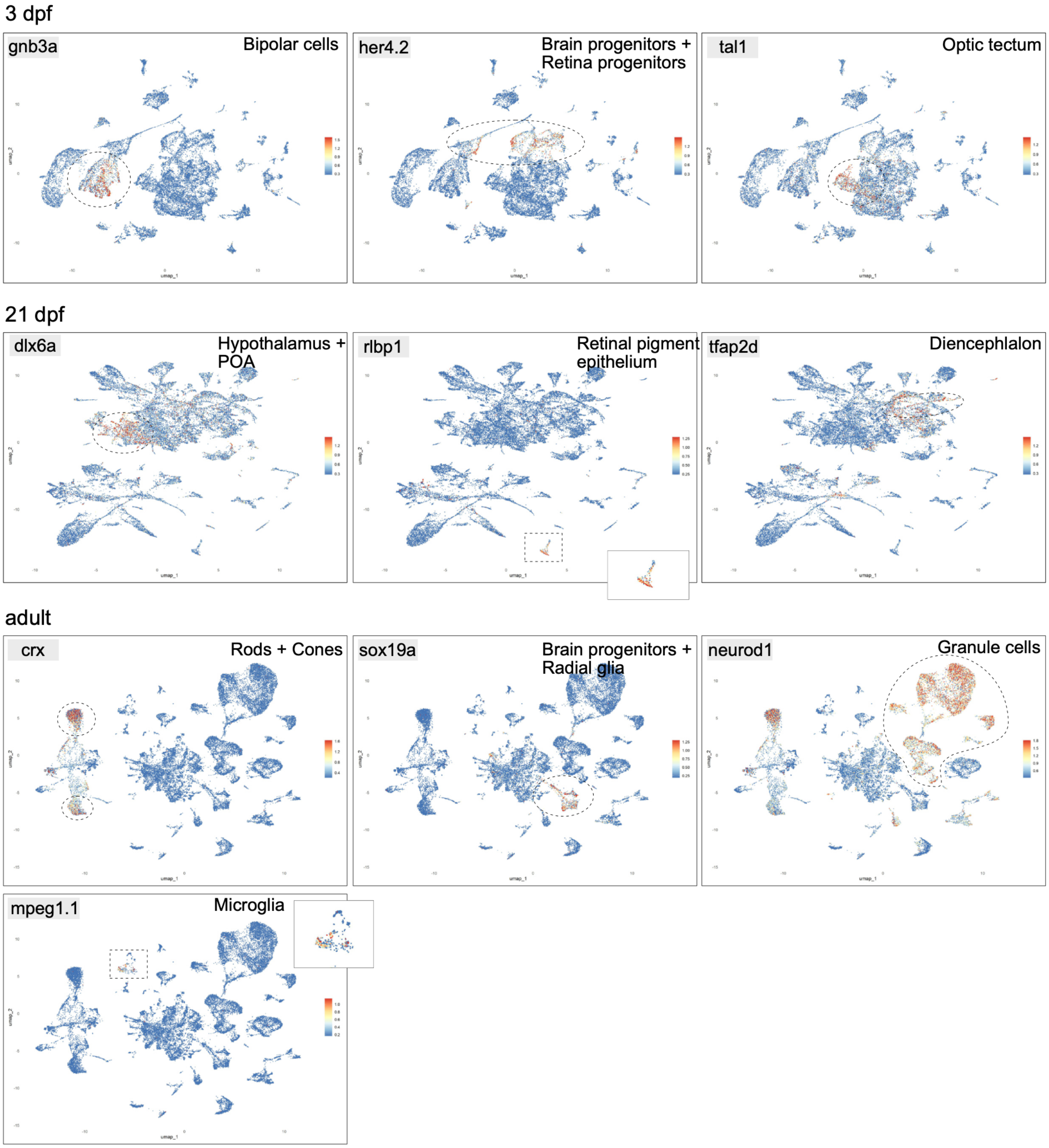


**Supplemental Fig. S1. Gene activity of canonical markers across brain and retinal cell populations.** Feature plots showing gene activity scores of canonical marker genes across major brain and retinal cell types at 3 dpf, 21 dpf, and adult stages.

A

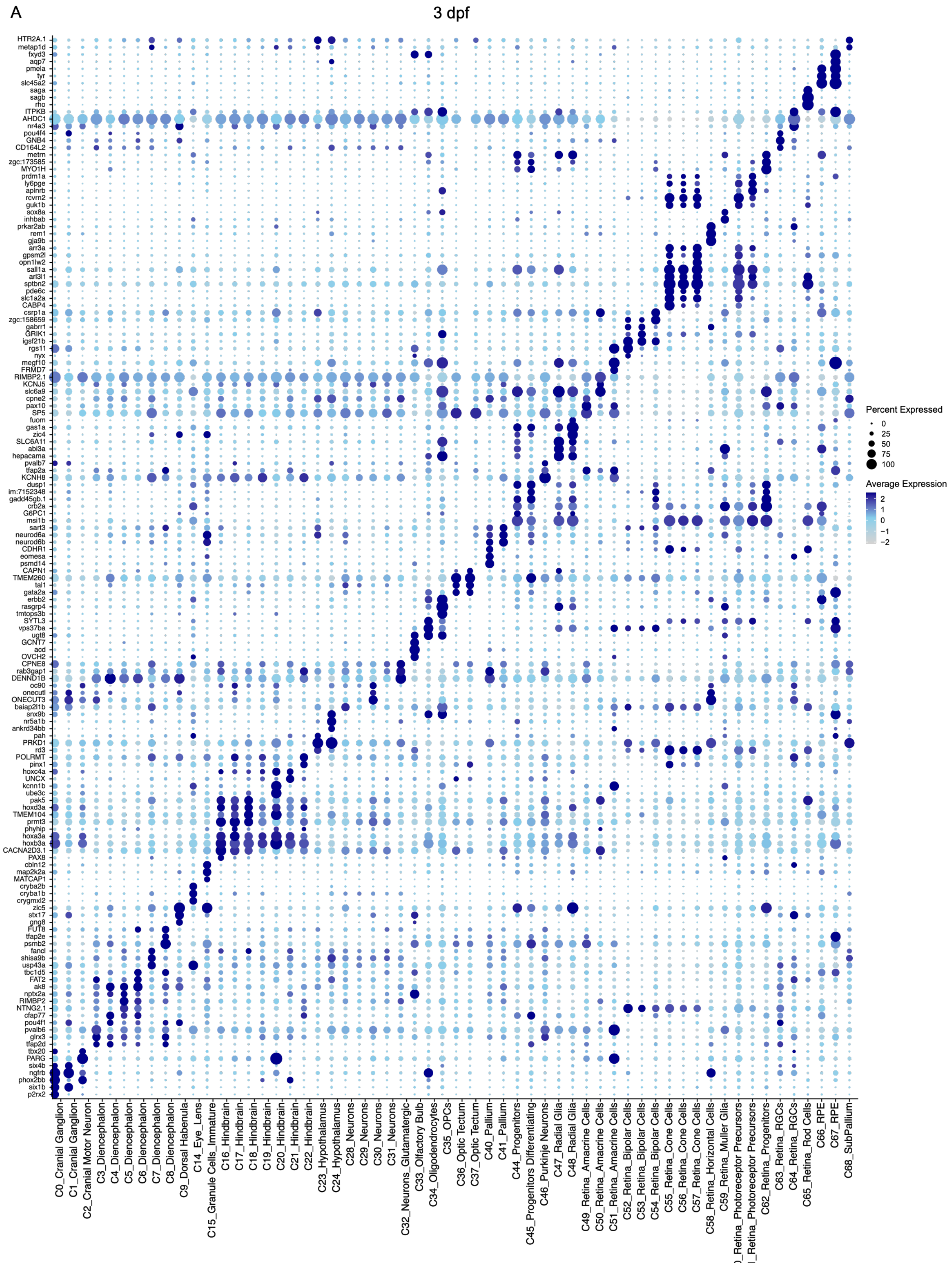
3 dpf
HTR2A.1
metap1d
fxyd3
aqp7
pmela
tyr
slc45a2
saga
sagb
rho
ITPKB
AHDC1
nr4a3
pou4f4
GNB4
CD164L2
metrn
zgc:173585
MYO1H
prdm1a
ly6pge
aplnrb
rcvrn2
guk1b
sox8a
inhbab
prkar2ab
rem1
gja9b
arr3a
gpsm2l
opn1lw2
sall1a
arl3l1
sptbn2
pde6c
slc1a2a
CABP4
csrp1a
zgc:158659
gabrr1
GRIK1
igsf21b
rgs11
nyx
megf10
FRMD7
RIMBP2.1
KCNJ5
slc6a9
cpne2
pax10
SP5
fuom
gas1a
zic4
SLC6A11
abi3a
hepacama
pvalb7
tfap2a
KCNH8
dusp1
im:7152348
gadd45gb.1
crb2a
G6PC1
msi1b
sart3
neurod6a
neurod6b
CDHR1
eomesa
psmd14
CAPN1
TMEM260
tal1
gata2a
erbb2
rasgrp4
tmtops3b
SYTL3
vps37ba
ugt8
GCNT7
acd
OVCH2
CPNE8
rab3gap1
DENND1B
oc90
onecutl
ONECUT3
baiap2l1b
snx9b
nr5a1b
ankrd34bb
pah
PRKD1
rd3
POLRMT
pinx1
hoxc4a
UNCX
kcnn1b
ube3c
pak5
hoxd3a
TMEM104
prmt3
phyhip
hoxa3a
hoxb3a
CACNA2D3.1
PAX8
cbln12
map2k2a
MATCAP1
cryba2b
cryba1b
crygmxl2
zic5
stx17
gng8
FUT8
tfap2e
psmb2
fancl
shisa9b
usp43a
tbc1d5
FAT2
ak8
nptx2a
RIMBP2
NTNG2.1
cfap77
pou4f1
pvalb6
glrx3
tfap2d
tbx20
PARG
six4b
ngfrb
phox2bb
six1b
p2rx2
C0_Cranial Ganglion
C1_Cranial Ganglion
C2_Cranial Motor Neuron
C3_Diencephalon
C4_Diencephalon
C5_Diencephalon
C6_Diencephalon
C7_Diencephalon
C8_Diencephalon
C9_Dorsal Habenula
C14_Eye_Lens
C15_Granule Cells_Immature
C16_Hindbrain
C17_Hindbrain
C18_Hindbrain
C19_Hindbrain
C20_Hindbrain
C21_Hindbrain
C22_Hindbrain
C23_Hypothalamus
C24_Hypothalamus
C28_Neurons
C29_Neurons
C30_Neurons
C31_Neurons
C32_Neurons_Glutamatergic
C33_Olfactory Bulb
C34_Oligodendrocytes
C35_OPCs
C36_Optic Tectum
C37_Optic Tectum
C40_Pallium
C41_Pallium
C44_Progenitors
C45_Progenitors Differentiating
C46_Purkinje Neurons
C47_Radial Glia
C48_Radial Glia
C49_Retina_Amacrine Cells
C50_Retina_Amacrine Cells
C51_Retina_Amacrine Cells
C52_Retina_Bipolar Cells
C53_Retina_Bipolar Cells
C54_Retina_Bipolar Cells
C55_Retina_Cone Cells
C56_Retina_Cone Cells
C57_Retina_Cone Cells
C58_Retina_Horizontal Cells
C59_Retina_Muller Glia
C60_Retina_Photoreceptor Precursors
C61_Retina_Photoreceptor Precursors
C62_Retina_Progenitors
C63_Retina_RGCs
C64_Retina_RGCs
C65_Retina_Rod Cells
C66_RPE
C67_RPE
C68_SubPallium
Percent Expressed
0
25
50
75
100
Average Expression
2
1
0
−1
−2

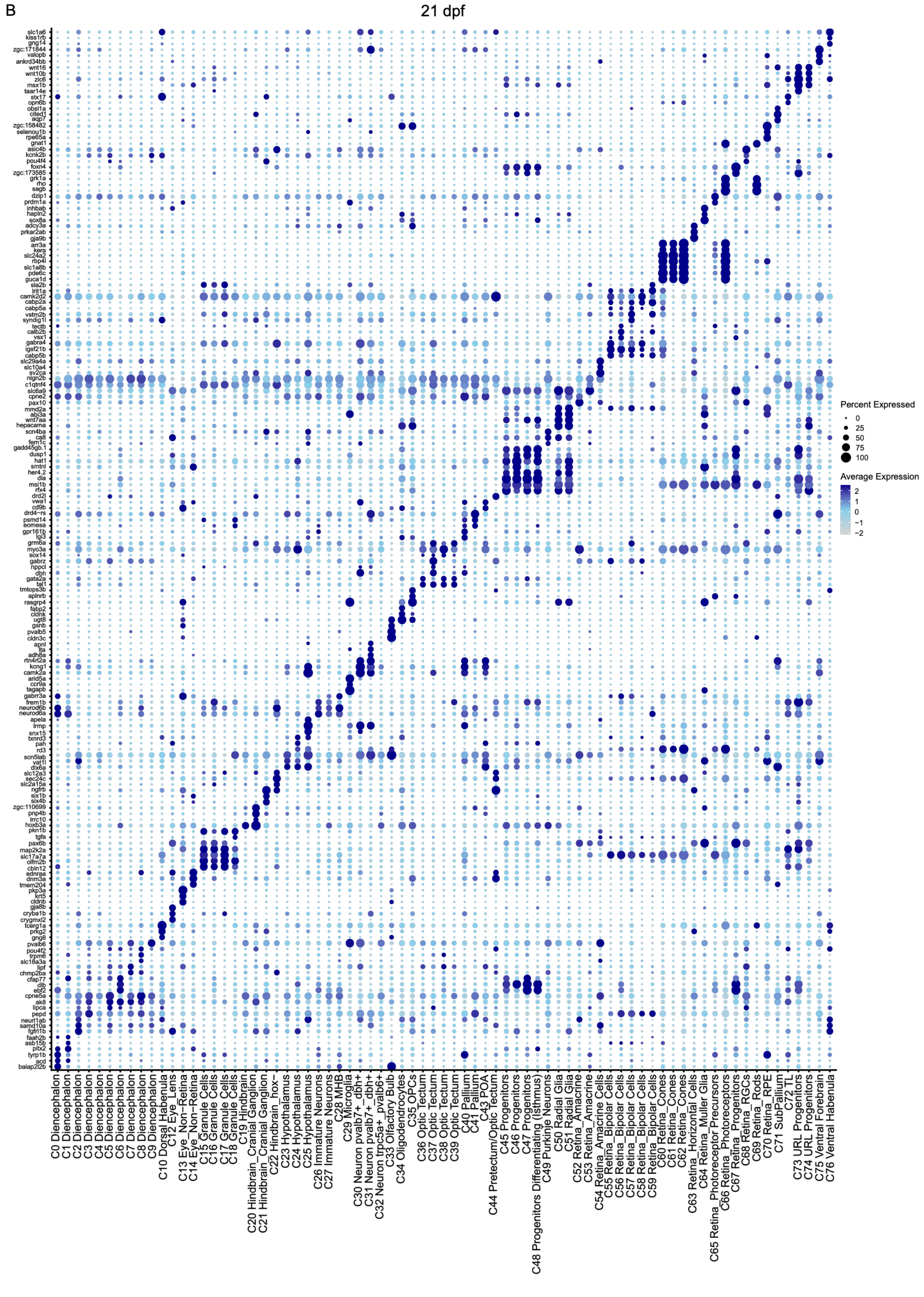
B
21 dpf
Percent Expressed
0
25
50
75
100
Average Expression
2
1
0
−1
−2
C0 Diencephalon
C1 Diencephalon
C2 Diencephalon
C3 Diencephalon
C4 Diencephalon
C5 Diencephalon
C6 Diencephalon
C7 Diencephalon
C8 Diencephalon
C9 Diencephalon
C10 Dorsal Habenula
C12 Eye_Lens
C13 Eye_Non−Retina
C14 Eye_Non−Retina
C15 Granule Cells
C16 Granule Cells
C17 Granule Cells
C18 Granule Cells
C19 Hindbrain
C20 Hindbrain_Cranial Ganglion
C21 Hindbrain_Cranial Ganglion
C22 Hindbrain_hox−
C23 Hypothalamus
C24 Hypothalamus
C25 Hypothalamus
C26 Immature Neurons
C27 Immature Neurons
C28 MHB
C29 Microglia
C30 Neuron pvalb7+_dbh+
C31 Neuron pvalb7+_dbh+
C32 Neuron trpc5a+_pvalb6+
C33 Olfactory Bulb
C34 Oligodendrocytes
C35 OPCs
C36 Optic Tectum
C37 Optic Tectum
C38 Optic Tectum
C39 Optic Tectum
C40 Pallium
C41 Pallium
C43 POA
C44 Pretectum/Optic Tectum
C45 Progenitors
C46 Progenitors
C47 Progenitors
C48 Progenitors Differentiating (Isthmus)
C49 Purkinje Neurons
C50 Radial Glia
C51 Radial Glia
C52 Retina_Amacrine
C53 Retina_Amacrine
C54 Retina_Amacrine Cells
C55 Retina_Bipolar Cells
C56 Retina_Bipolar Cells
C57 Retina_Bipolar Cells
C58 Retina_Bipolar Cells
C59 Retina_Bipolar Cells
C60 Retina_Cones
C61 Retina_Cones
C62 Retina_Cones
C63 Retina_Horizontal Cells
C64 Retina_Muller Glia
C65 Retina_Photoreceptor Precursors
C66 Retina_Photoreceptors
C67 Retina_Progenitors
C68 Retina_RGCs
C69 Retina_Rods
C70 Retina_RPE
C71 SubPallium
C72 TL
C73 URL Progenitors
C74 URL Progenitors
C75 Ventral Forebrain
C76 Ventral Habenula

C

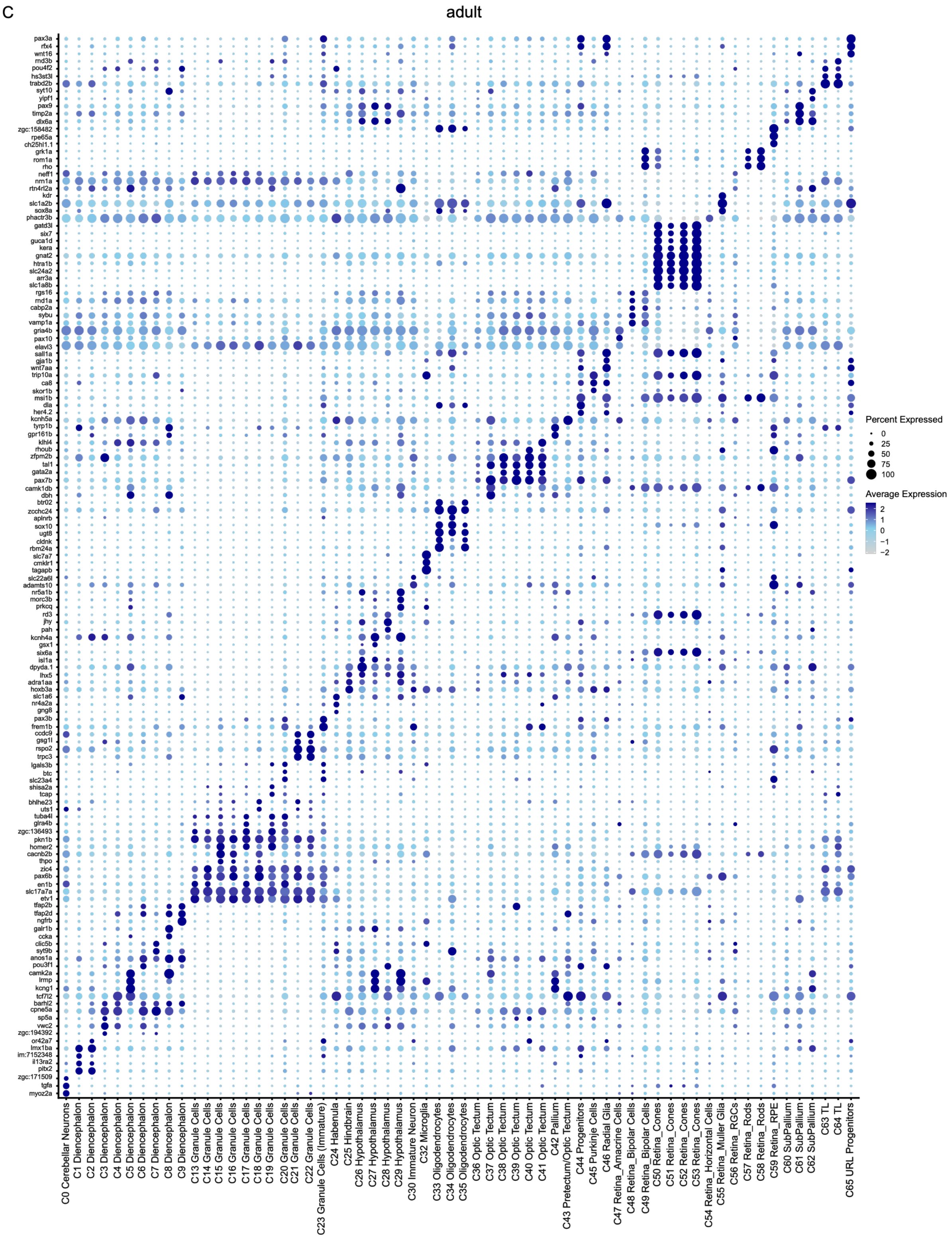


**Supplemental Fig. S2. Cluster-enriched gene activity signatures.** Dot plots of gene activity scores of select markers (rows) enriched in each cluster (columns) in the (A) 3 dpf, (B) 21 dpf, and (C) adult scATAC-seq datasets. Dot size indicates the percentage of cells expressing the marker; color represents the average scaled expression level.

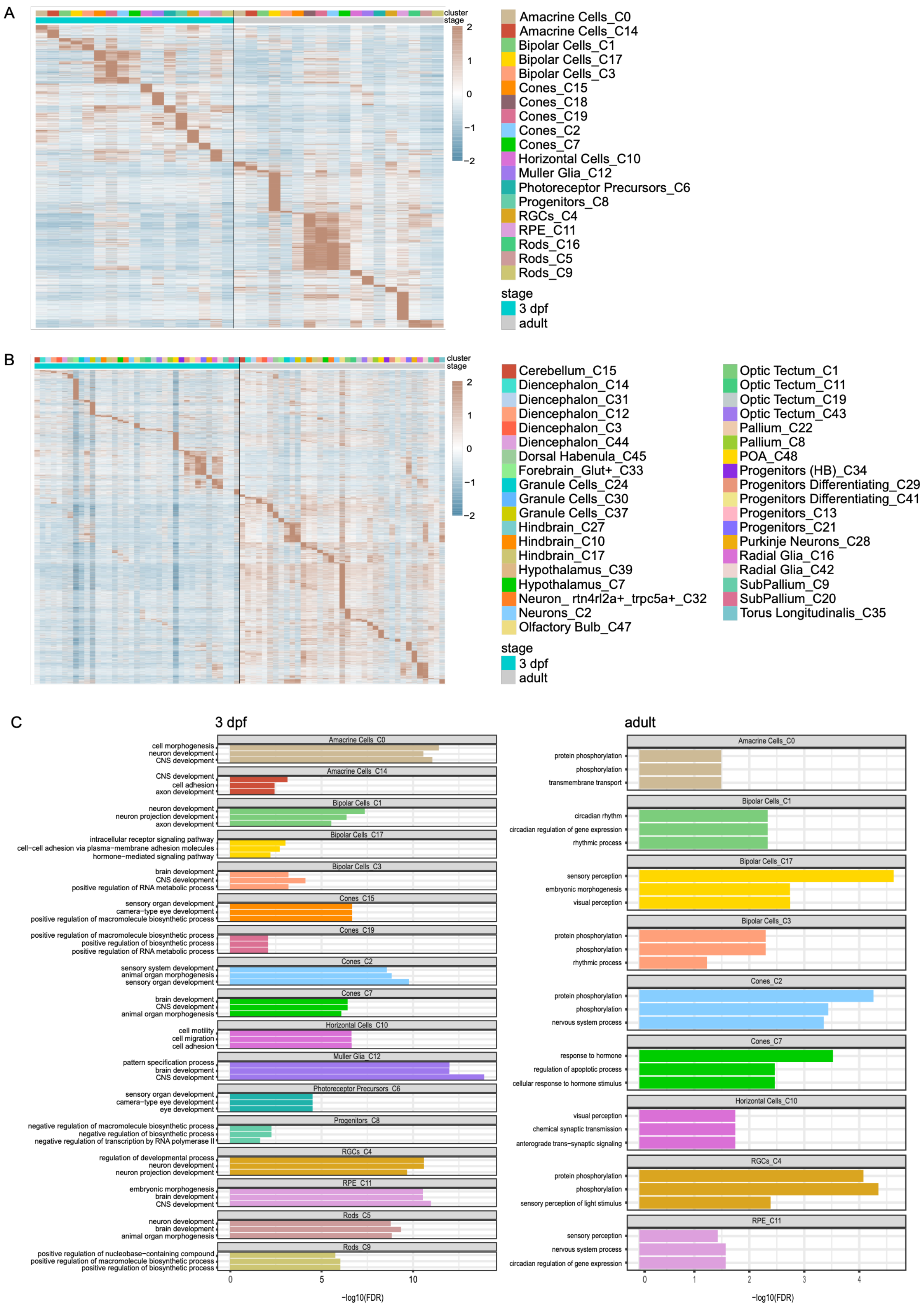
A
cluster
stage
2
1
0
−1
−2
Amacrine Cells_C0
Amacrine Cells_C14
Bipolar Cells_C1
Bipolar Cells_C17
Bipolar Cells_C3
Cones_C15
Cones_C18
Cones_C19
Cones_C2
Cones_C7
Horizontal Cells_C10
Muller Glia_C12
Photoreceptor Precursors_C6
Progenitors_C8
RGCs_C4
RPE_C11
Rods_C16
Rods_C5
Rods_C9
stage
3 dpf
adult
B
cluster
stage
2
1
0
−1
−2
Cerebellum_C15
Diencephalon_C14
Diencephalon_C31
Diencephalon_C12
Diencephalon_C3
Diencephalon_C44
Dorsal Habenula_C45
Forebrain_Glut+_C33
Granule Cells_C24
Granule Cells_C30
Granule Cells_C37
Hindbrain_C27
Hindbrain_C10
Hindbrain_C17
Hypothalamus_C39
Hypothalamus_C7
Neuron_ rtn4rl2a+_trpc5a+_C32
Neurons_C2
Olfactory Bulb_C47
Optic Tectum_C1
Optic Tectum_C11
Optic Tectum_C19
Optic Tectum_C43
Pallium_C22
Pallium_C8
POA_C48
Progenitors (HB)_C34
Progenitors Differentiating_C29
Progenitors Differentiating_C41
Progenitors_C13
Progenitors_C21
Purkinje Neurons_C28
Radial Glia_C16
Radial Glia_C42
SubPallium_C9
SubPallium_C20
Torus Longitudinalis_C35
stage
3 dpf
adult
C
3 dpf
Amacrine Cells_C0
cell morphogenesis
neuron development
CNS development
Amacrine Cells_C14
CNS development
cell adhesion
axon development
Bipolar Cells_C1
neuron development
neuron projection development
axon development
Bipolar Cells_C17
intracellular receptor signaling pathway
cell−cell adhesion via plasma−membrane adhesion molecules
hormone−mediated signaling pathway
Bipolar Cells_C3
brain development
CNS development
positive regulation of RNA metabolic process
Cones_C15
sensory organ development
camera−type eye development
positive regulation of macromolecule biosynthetic process
Cones_C19
positive regulation of macromolecule biosynthetic process
positive regulation of biosynthetic process
positive regulation of RNA metabolic process
Cones_C2
sensory system development
animal organ morphogenesis
sensory organ development
Cones_C7
brain development
CNS development
animal organ morphogenesis
Horizontal Cells_C10
cell motility
cell migration
cell adhesion
Muller Glia_C12
pattern specification process
brain development
CNS development
Photoreceptor Precursors_C6
sensory organ development
camera−type eye development
eye development
Progenitors_C8
negative regulation of macromolecule biosynthetic process
negative regulation of biosynthetic process
negative regulation of transcription by RNA polymerase II
RGCs_C4
regulation of developmental process
neuron development
neuron projection development
RPE_C11
embryonic morphogenesis
brain development
CNS development
Rods_C5
neuron development
brain development
animal organ morphogenesis
Rods_C9
positive regulation of nucleobase−containing compound
positive regulation of macromolecule biosynthetic process
positive regulation of biosynthetic process
0
5
10
−log10(FDR)
adult
Amacrine Cells_C0
protein phosphorylation
phosphorylation
transmembrane transport
Bipolar Cells_C1
circadian rhythm
circadian regulation of gene expression
rhythmic process
Bipolar Cells_C17
sensory perception
embryonic morphogenesis
visual perception
Bipolar Cells_C3
protein phosphorylation
phosphorylation
rhythmic process
Cones_C2
protein phosphorylation
phosphorylation
nervous system process
Cones_C7
response to hormone
regulation of apoptotic process
cellular response to hormone stimulus
Horizontal Cells_C10
visual perception
chemical synaptic transmission
anterograde trans−synaptic signaling
RGCs_C4
protein phosphorylation
phosphorylation
sensory perception of light stimulus
RPE_C11
sensory perception
nervous system process
circadian regulation of gene expression
0
1
2
3
4
−log10(FDR)

**Supplemental Fig. S3: Biological programs linked to differentially accessible peaks.** (A–B) Heatmaps showing accessibility of all DA peaks (rows) enriched at 3 dpf or adult stages within cell types (columns) of the (A) retina and (B) brain. Mean fragment counts per timepoint were computed using AggregateExpression, log transformed, and z-score normalized across timepoints for visualization. (C) GO enrichment analysis of genes linked to DA peaks for each retina cluster at 3 dpf and adult stages. The top three GO terms per cluster based on adjusted p-value are shown. The x-axis indicates −log10(FDR).

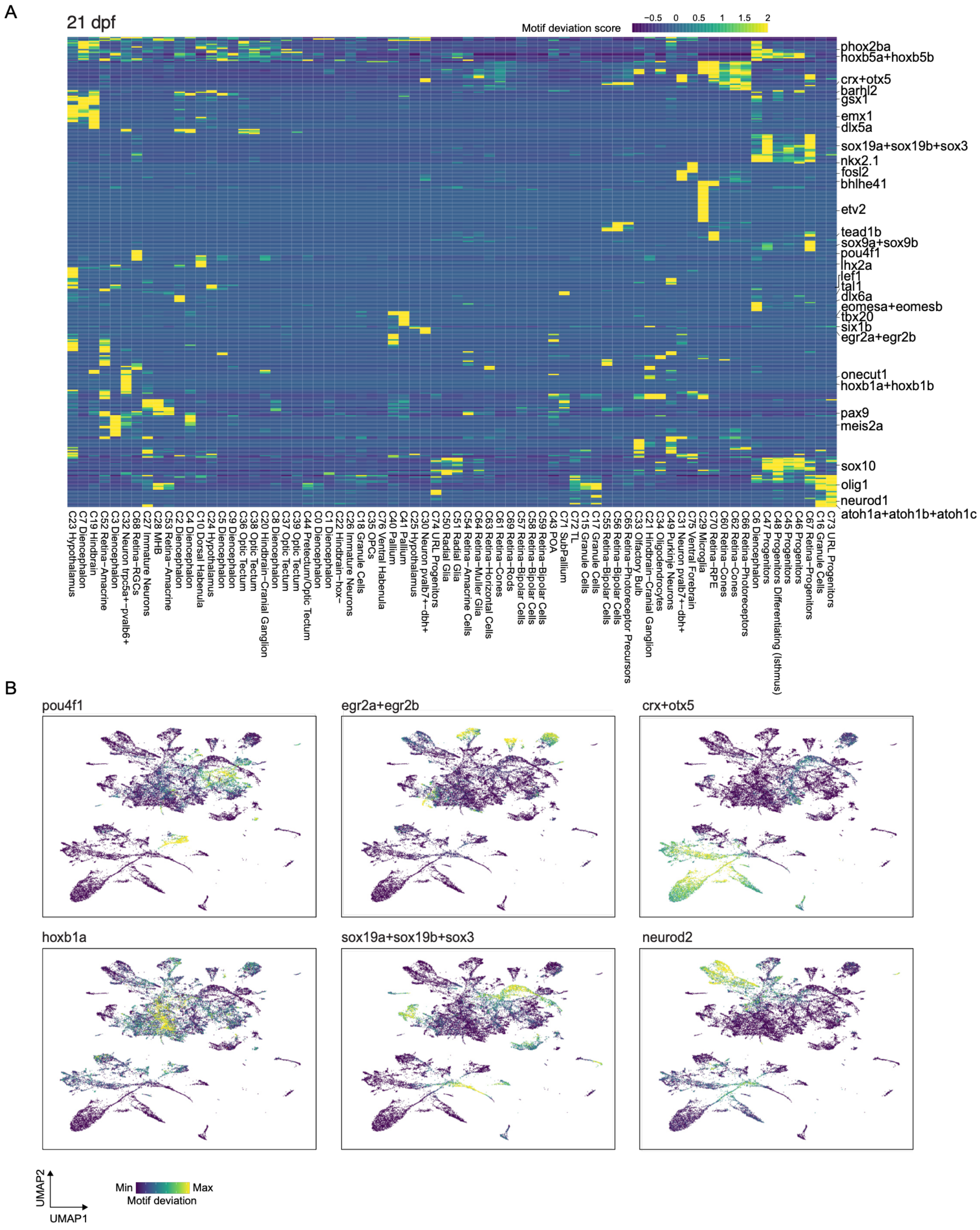


**Supplemental Fig. S4. Transcription factor motif accessibility in 21 dpf brain and retina.** (A) Heatmap of chromVAR TF motif deviation scores across 21 dpf clusters. (B) UMAP projections of 21 dpf scATAC-seq data colored by chromVAR motif deviation scores for selected TFs.

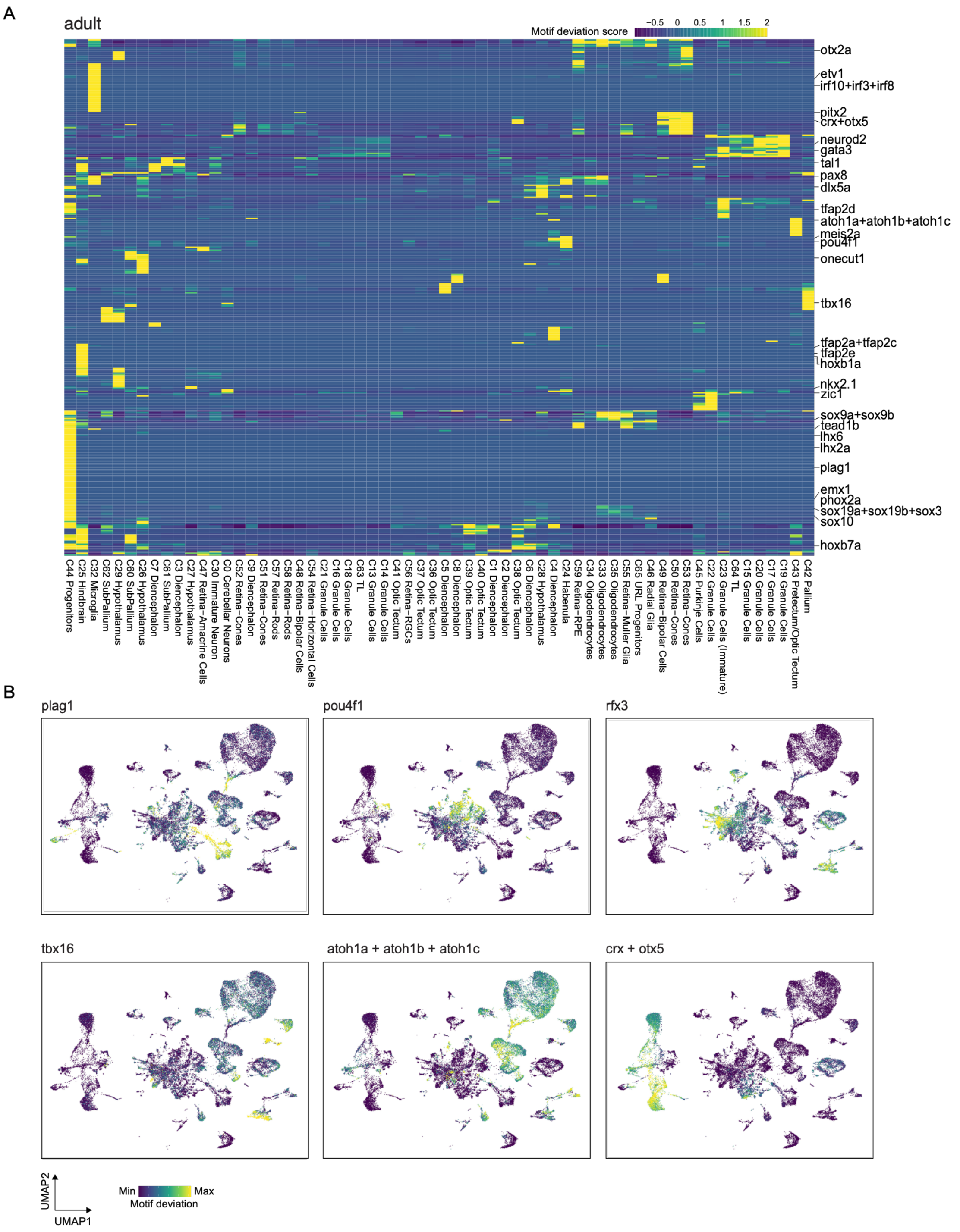


**Supplemental Fig. S5. Transcription factor motif accessibility in adult brain and retina.** (A) Heatmap of chromVAR TF motif deviation scores across adult clusters. (B) UMAP projections of adult scATAC-seq data colored by chromVAR motif deviation scores for selected TFs.

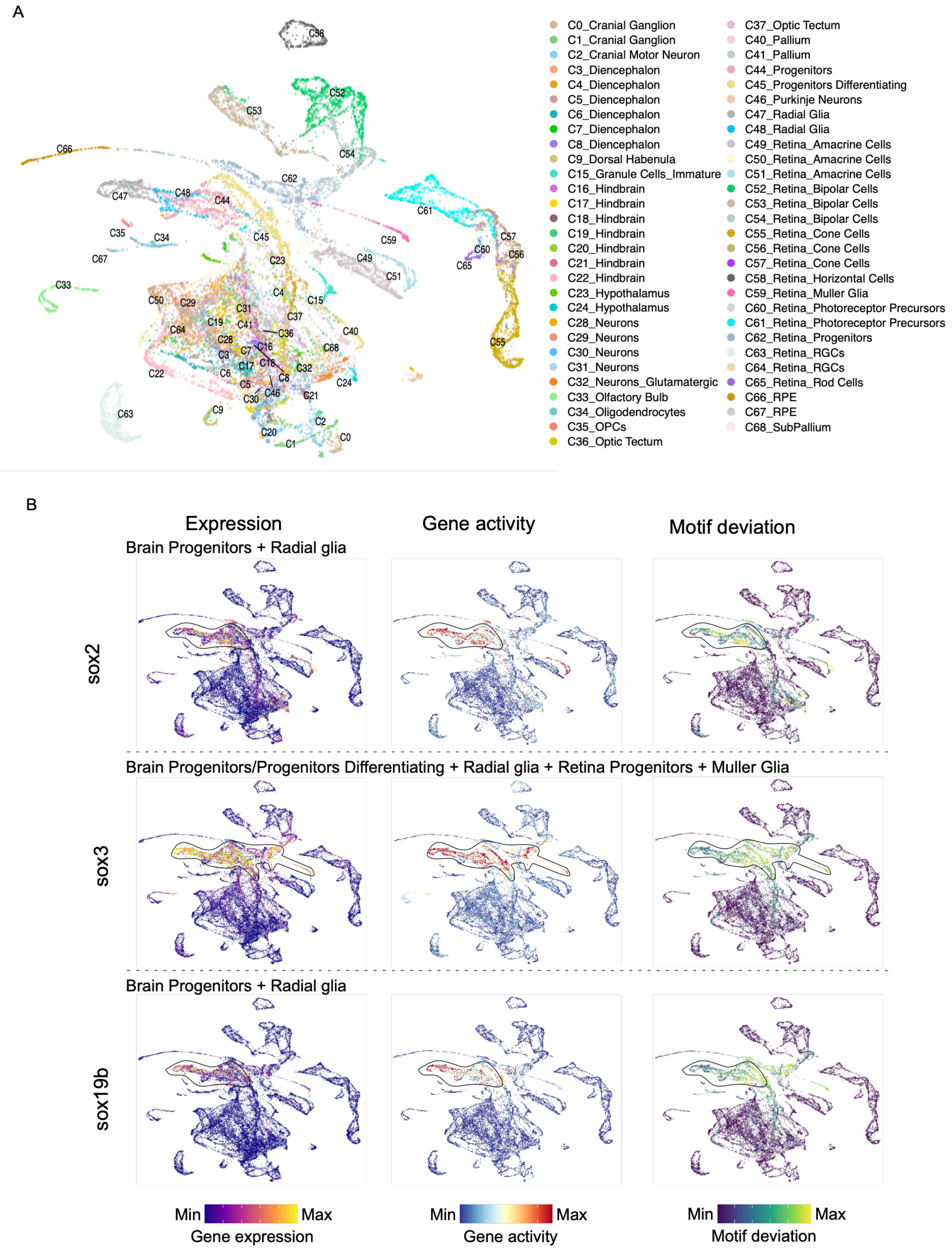


**Supplemental Fig. S6**. **Integrated scATAC-seq and scRNA-seq 3 dpf dataset.** (A) UMAP projection of the integrated 3 dpf scATAC-scRNA dataset colored by cell type. Only cells from the scATAC-seq experiment are shown. (B) UMAP projections colored by motif accessibility, gene activity, and gene expression for SoxB1 family members.

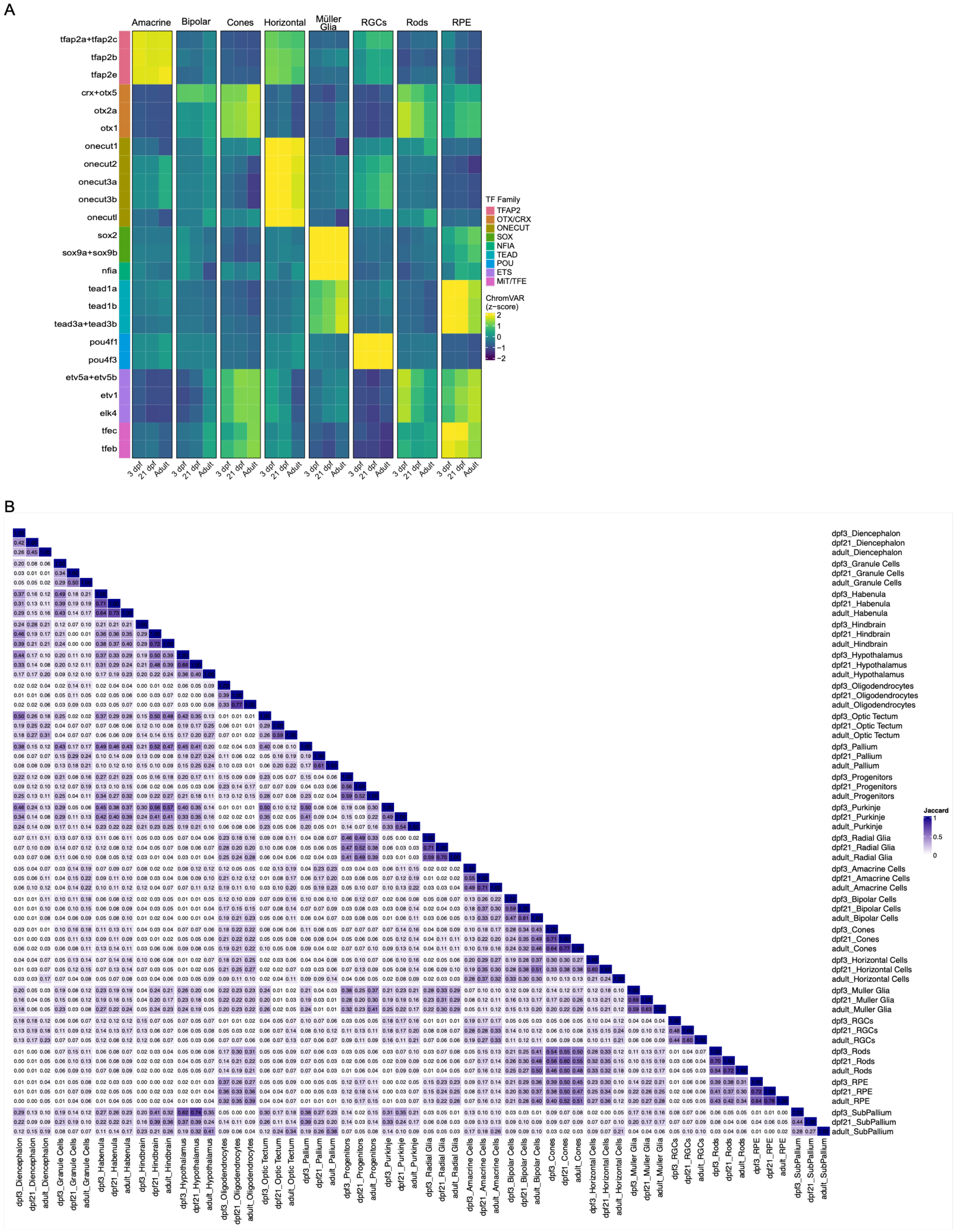


**Supplemental Fig. S7. Transcription factor motif similarity by cell type and region.** (A) Heatmap of chromVAR motif accessibility (z-scored deviation scores) for selected TF families across major retinal cell classes across timepoints. (B) Similarity of aggregate chromVAR-derived motif accessibility profiles within indicated cell types and regions across timepoints. Heatmap shows Jaccard index overlap.

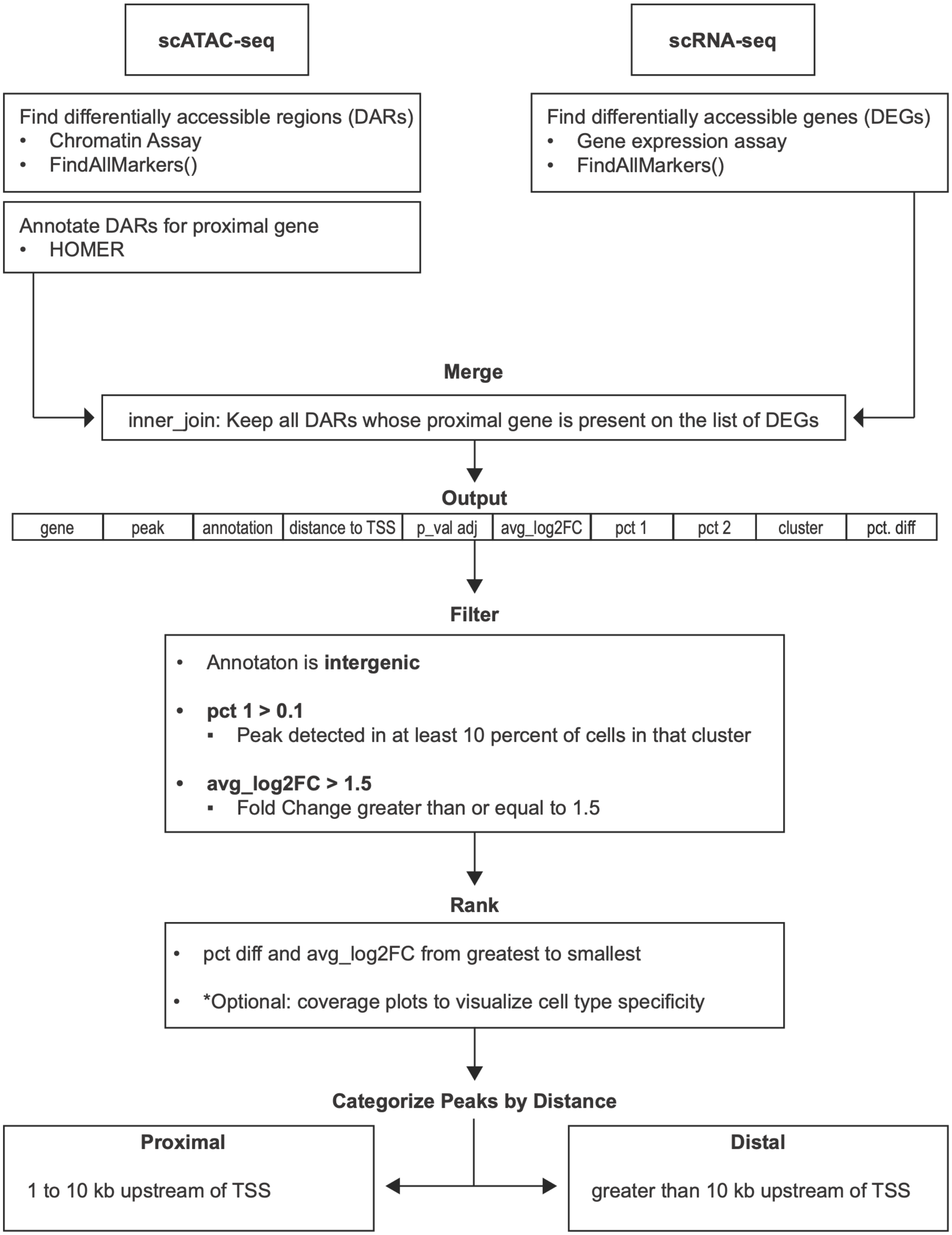


**Supplemental Fig. S8. Bioinformatic pipeline for prioritization of cis-regulatory elements for testing.** Flowchart schematic outlining the computational workflow used to identify candidate CREs associated with differentially expressed genes. Differentially accessible regions (DARs) identified from 3 dpf scATAC-seq were annotated to proximal genes and intersected with differentially expressed genes (DEGs) from 3 dpf scRNA-seq. Peaks were filtered based on genomic annotation, accessibility thresholds, and fold-change criteria, ranked by accessibility metrics, and categorized by distance to transcription start sites to define proximal and distal candidate regulatory elements.

A

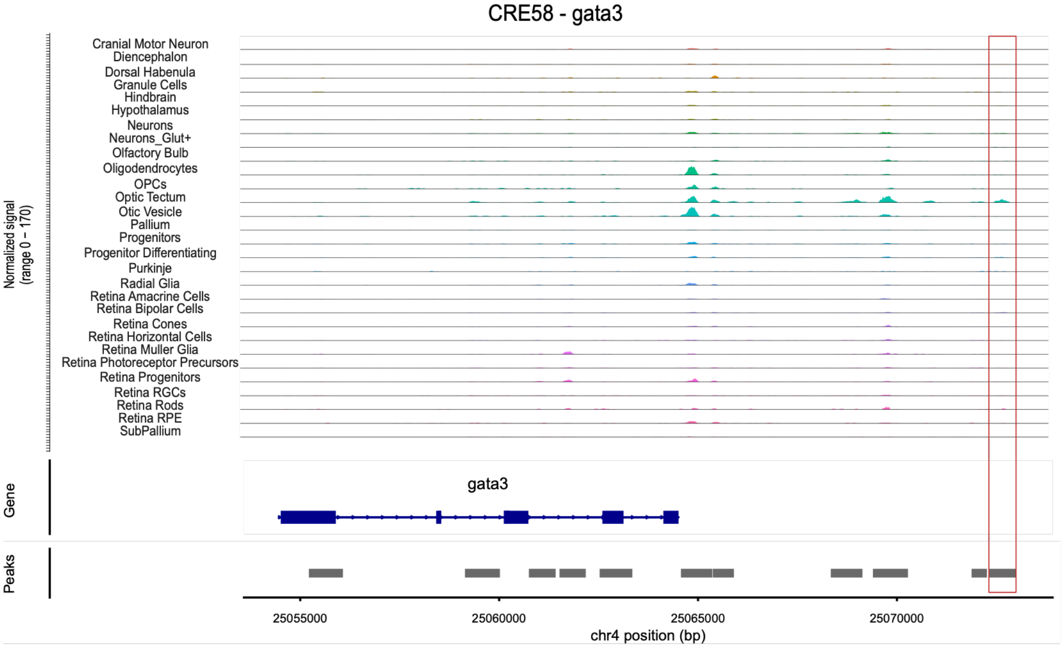
CRE58 - gata3
Normalized signal (range 0 – 170)
Cranial Motor Neuron
Diencephalon
Dorsal Habenula
Granule Cells
Hindbrain
Hypothalamus
Neurons
Neurons_Glut+
Olfactory Bulb
Oligodendrocytes
OPCs
Optic Tectum
Otic Vesicle
Pallium
Progenitors
Progenitor Differentiating
Purkinje
Radial Glia
Retina Amacrine Cells
Retina Bipolar Cells
Retina Cones
Retina Horizontal Cells
Retina Muller Glia
Retina Photoreceptor Precursors
Retina Progenitors
Retina RGCs
Retina Rods
Retina RPE
SubPallium
Gene
gata3
Peaks
25055000
25060000
25065000
25070000
chr4 position (bp)

B

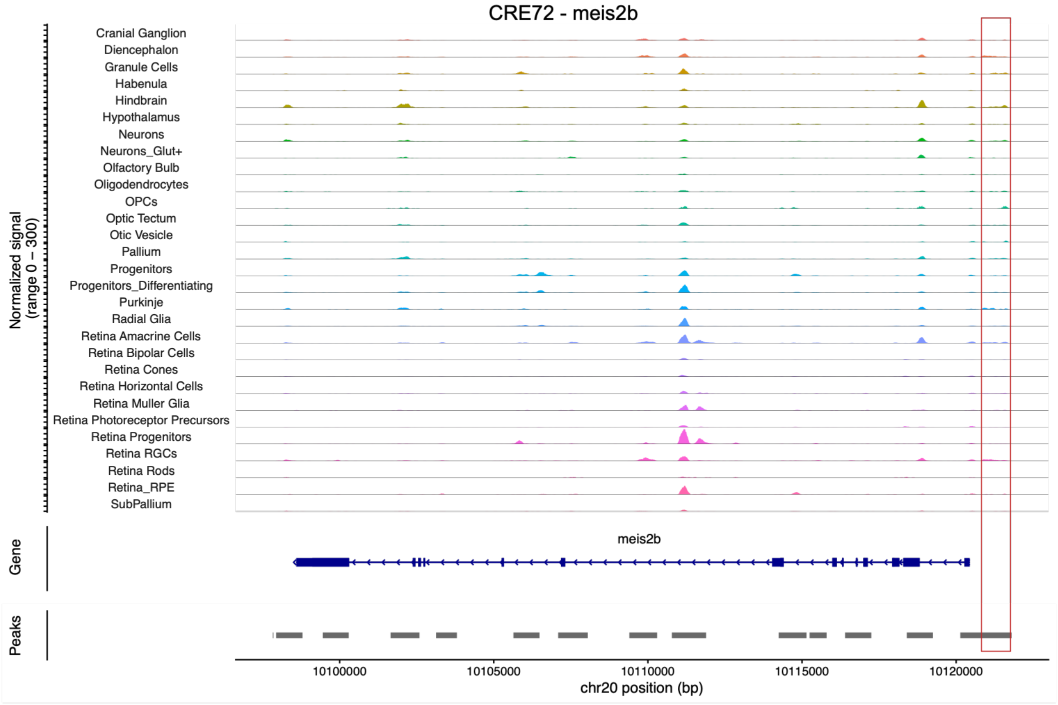
CRE72 - meis2b
Normalized signal (range 0 – 300)
Cranial Ganglion
Diencephalon
Granule Cells
Habenula
Hindbrain
Hypothalamus
Neurons
Neurons_Glut+
Olfactory Bulb
Oligodendrocytes
OPCs
Optic Tectum
Otic Vesicle
Pallium
Progenitors
Progenitors_Differentiating
Purkinje
Radial Glia
Retina Amacrine Cells
Retina Bipolar Cells
Retina Cones
Retina Horizontal Cells
Retina Muller Glia
Retina Photoreceptor Precursors
Retina Progenitors
Retina RGCs
Retina Rods
Retina_RPE
SubPallium
Gene
meis2b
Peaks
10100000
10105000
10110000
10115000
10120000
chr20 position (bp)

C

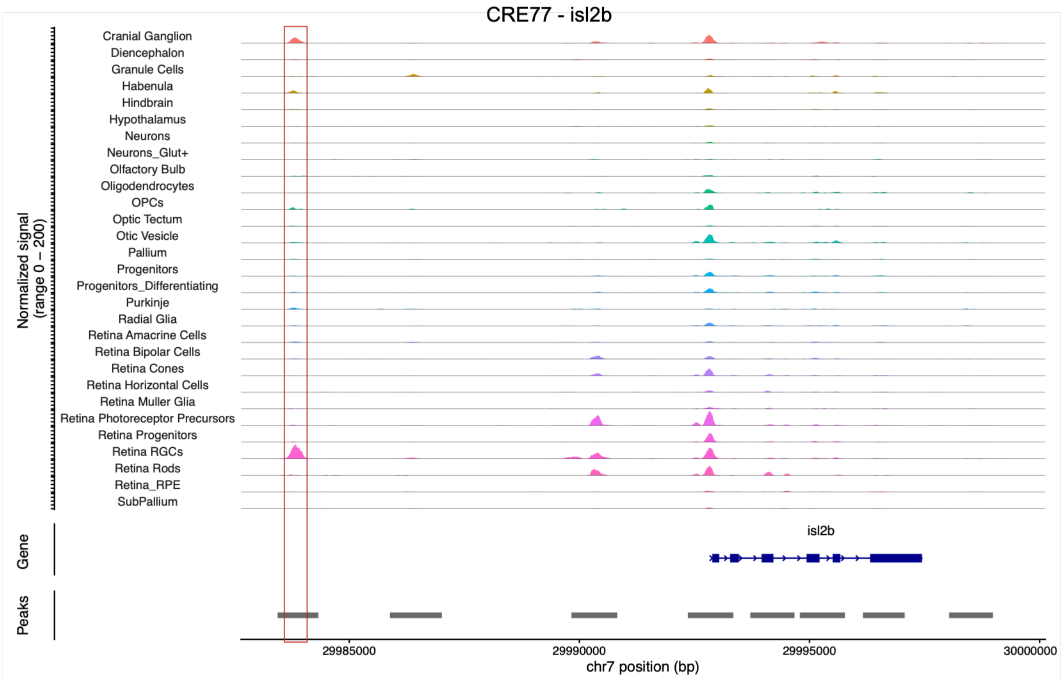


D

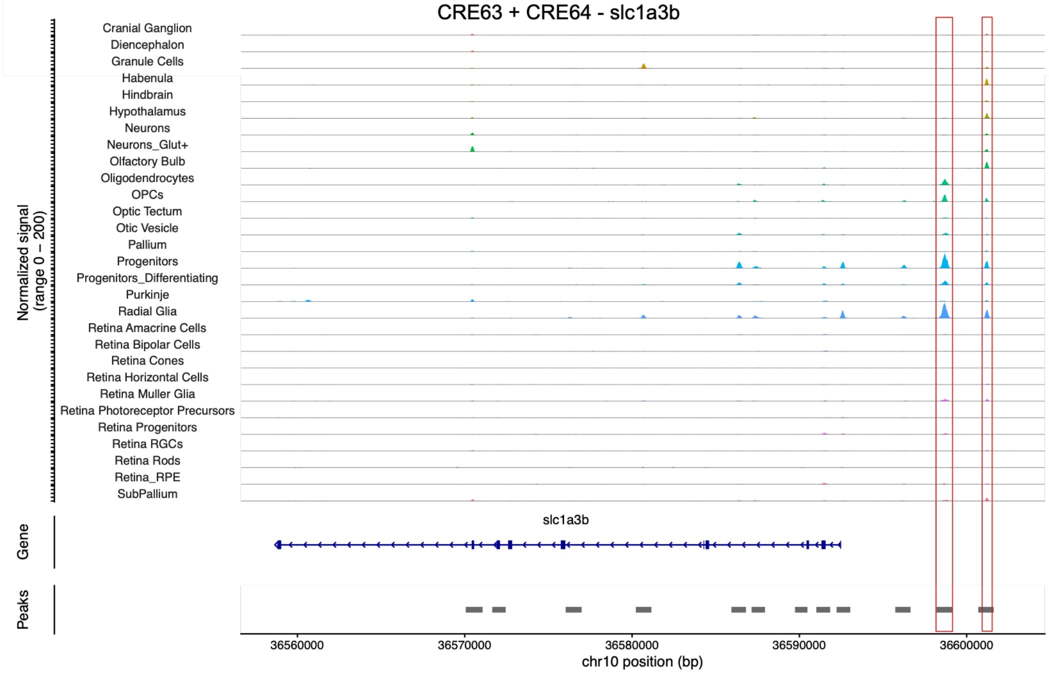


**Supplemental Fig S9. Genome coverage plots of cis-regulatory elements tested for enhancer activity.** (A-D) Genome browser views showing normalized scATAC-seq signal across representative loci for CREs selected for functional validation. Accessibility profiles are displayed across subsets of annotated cell types, with the nearby associated gene indicated below each locus. Red boxes highlight the genomic location of the candidate CRE tested in vivo. Coordinates are shown along the x-axis, and normalized accessibility signal is shown on the y-axis.

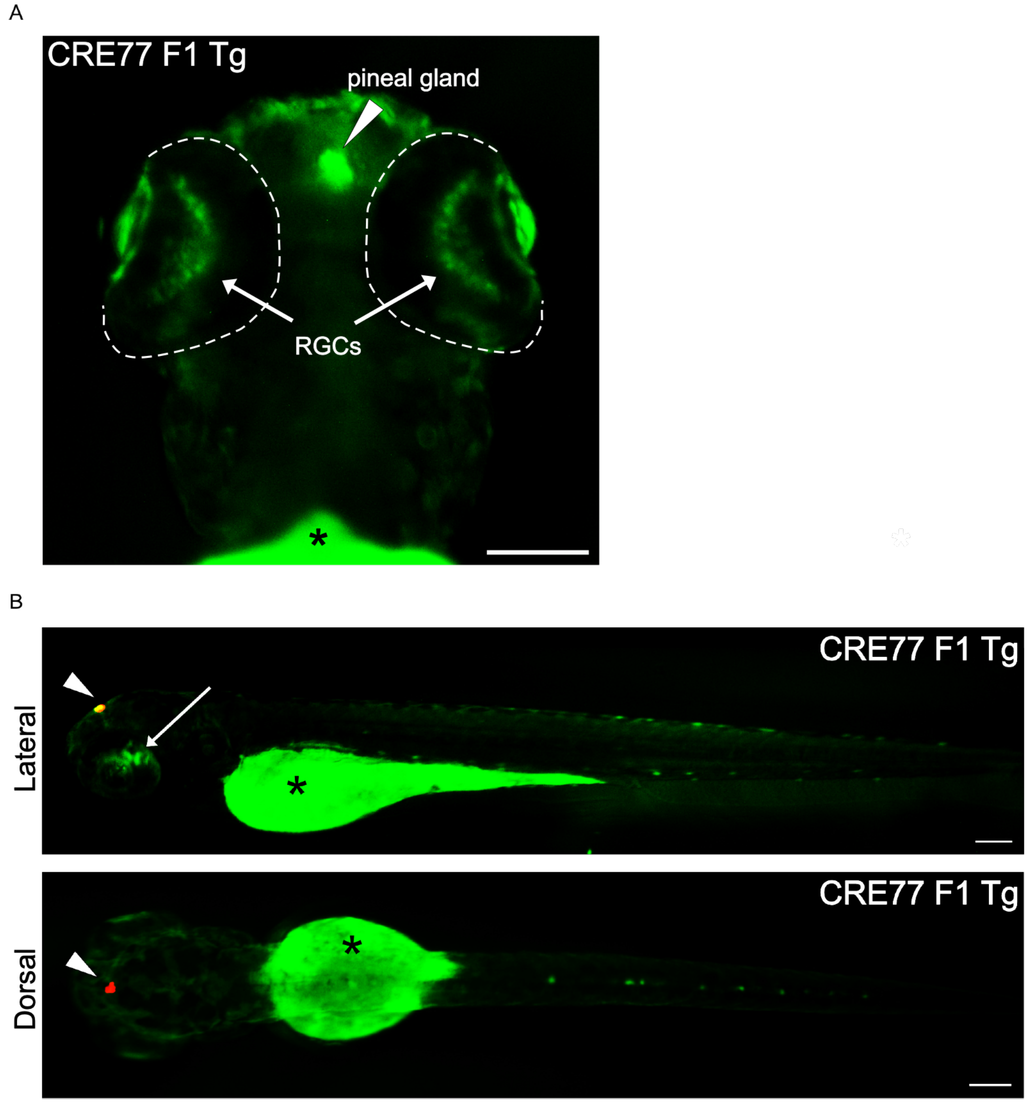


**Supplemental Fig. S10. Establishment of CRE77 F1 transgenics.** (A) Representative fluorescence image of F1 animals generated from a CRE77 F0 founder. Eyes are outlined in white dashed lines. White arrows indicate eGFP signal in the retina corresponding to RGCs, and the white arrowhead indicates the pineal gland. Strong mCherry fluorescence in the pineal gland bleeds through into the GFP channel. (B) Representative whole-body fluorescence images of CRE77 F1 animals. Scale bars, 100 µm. Asterisk, autofluorescence in yolk.

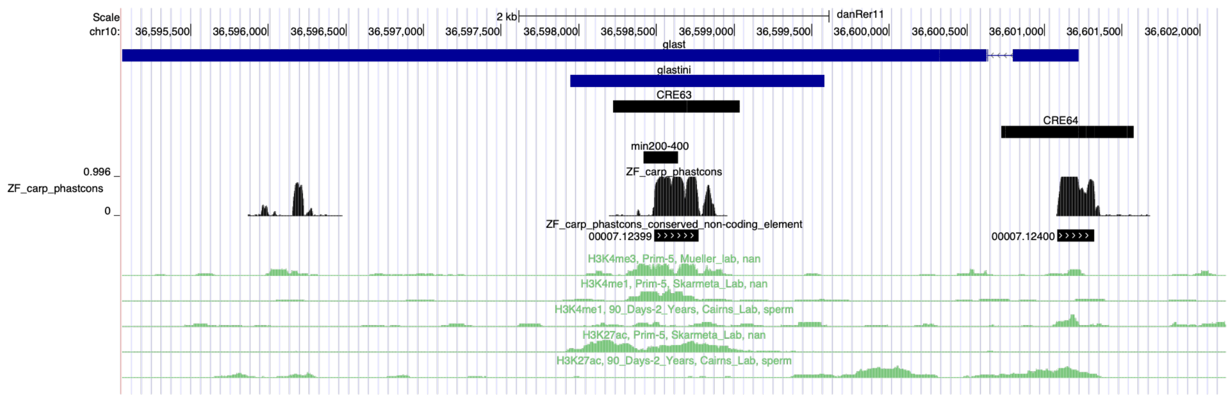


**Supplemental Fig. S11. Evolutionary conservation of the *slc1a3b* locus.** Genome browser view of the *slc1a3b* locus (danRer11; chr10) showing the positions of CRE63 and CRE64 relative to 9.5kb (glast) and 3 kb (glastini) promoters. Tracks display publicly available ChIP–seq datasets for H3K27ac, H3K4me1, H3K4me3, and H3K27me3 from embryonic (Prim- 5) and adult tissues, as indicated. PhastCons conservation scores across teleost species (ZF-carp alignment) are shown below, with conserved non-coding elements highlighted. CRE63 and CRE64 overlap regions of high evolutionary conservation. Position of the min200-400 fragment is indicated.

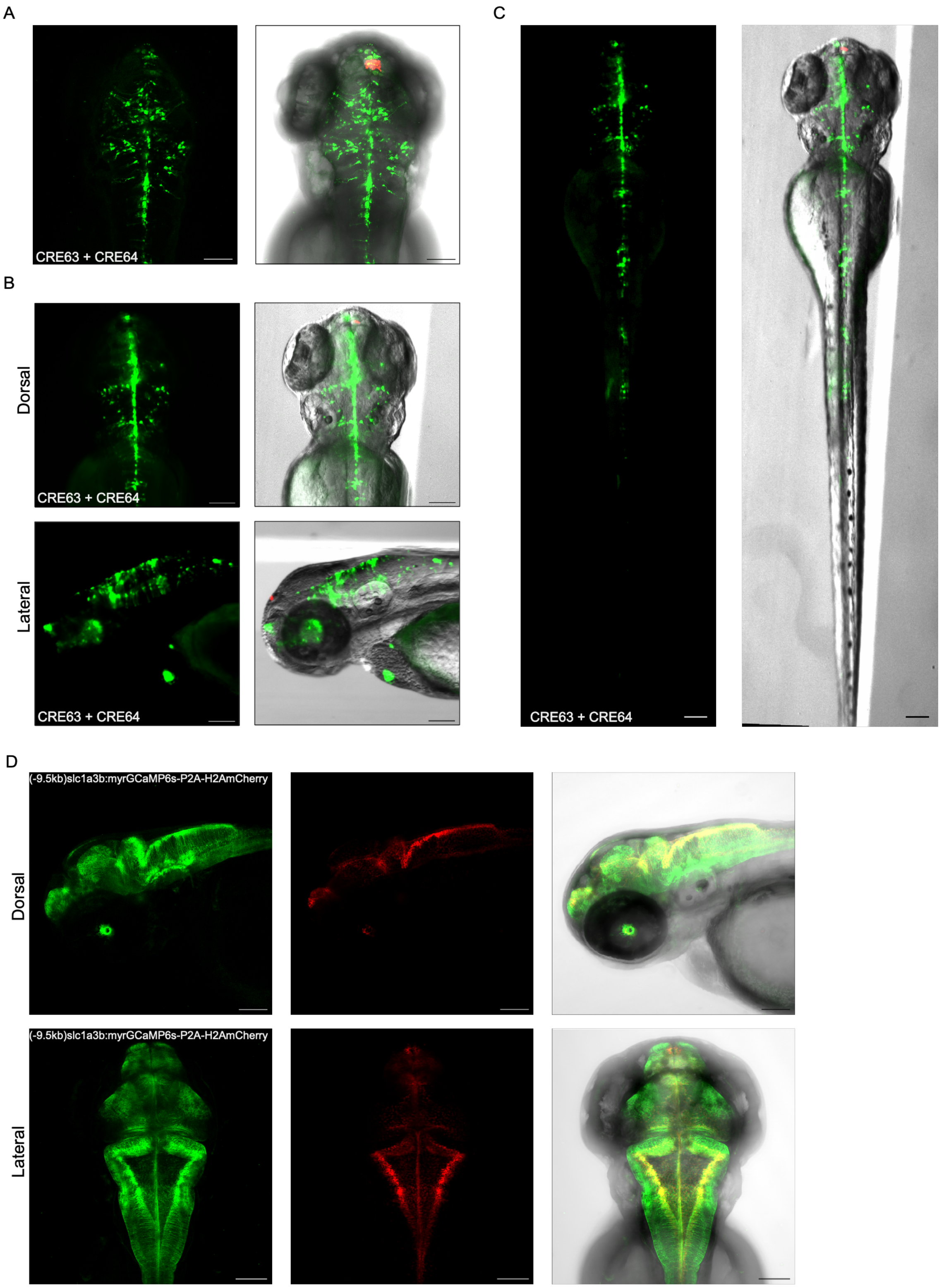


**Supplemental Fig. S12. Characterization of CRE63+CRE64 reporter activity.** (A-C) Representative images of 3 dpf zebrafish showing mosaic expression of the concatenated CRE63+CRE64 reporter construct. The pineal gland is marked by mCherry (red). (A) Maximum-intensity projections of confocal images of the head showing robust radial glia-associated eGFP expression in the brain. (B) Fluorescent images of the head with eGFP signal detected in the brain. (C) Whole-body fluorescent images with eGFP detected in the brain and spinal cord. (D) Confocal images of a previously published (-9.5kb)*slc1a3b*:myrGCaMP6s-P2A-H2AmCherry transgenic line (Chen et al. 2020). Cell bodies are labeled in red (H2A-mCherry) and cellular projections in green (myrGCaMP6s), highlighting radial glial morphology in the brain. Scale bars, 100 µm.

A

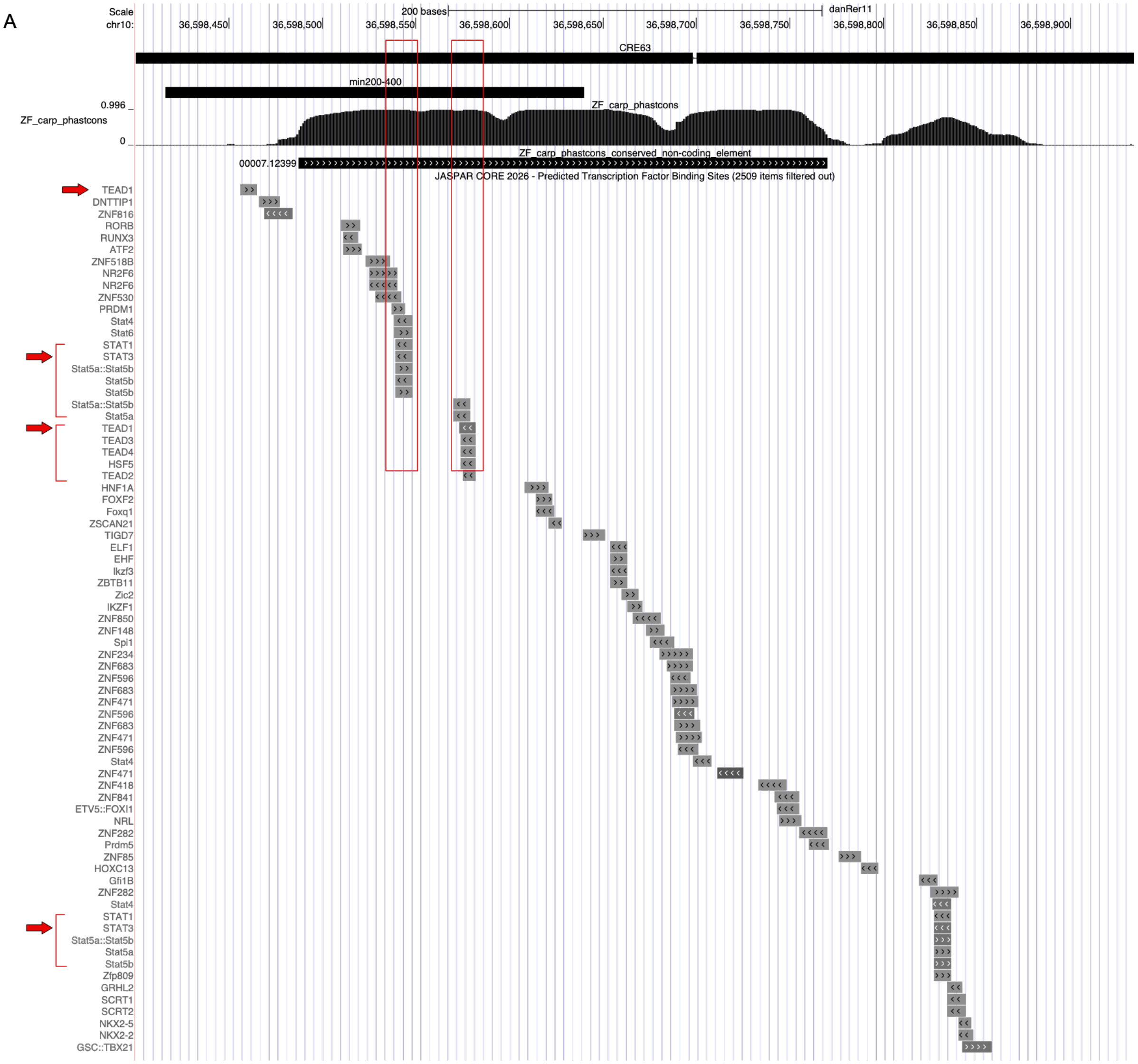

Scale
chr10:
200 bases
danRer11
36,598,450
36,598,500
36,598,550
36,598,600
36,598,650
36,598,700
36,598,750
36,598,800
36,598,850
36,598,900
CRE63
min200-400
ZF_carp_phastcons
0.996
0
ZF_carp_phastcons_conserved_non-coding_element
00007.12399
JASPAR CORE 2026 - Predicted Transcription Factor Binding Sites (2509 items filtered out)
TEAD1
DNTTIP1
ZNF816
RORB
RUNX3
ATF2
ZNF518B
NR2F6
NR2F6
ZNF530
PRDM1
Stat4
Stat6
STAT1
STAT3
Stat5a::Stat5b
Stat5b
Stat5b
Stat5a::Stat5b
Stat5a
TEAD1
TEAD3
TEAD4
HSF5
TEAD2
HNF1A
FOXF2
Foxq1
ZSCAN21
TIGD7
ELF1
EHF
Ikzf3
ZBTB11
Zic2
IKZF1
ZNF850
ZNF148
Spi1
ZNF234
ZNF683
ZNF596
ZNF683
ZNF471
ZNF596
ZNF683
ZNF471
ZNF596
Stat4
ZNF471
ZNF418
ZNF841
ETV5::FOXI1
NRL
ZNF282
Prdm5
ZNF85
HOXC13
Gfi1B
ZNF282
Stat4
STAT1
STAT3
Stat5a::Stat5b
Stat5a
Stat5b
Zfp809
GRHL2
SCRT1
SCRT2
NKX2-5
NKX2-2
GSC::TBX21

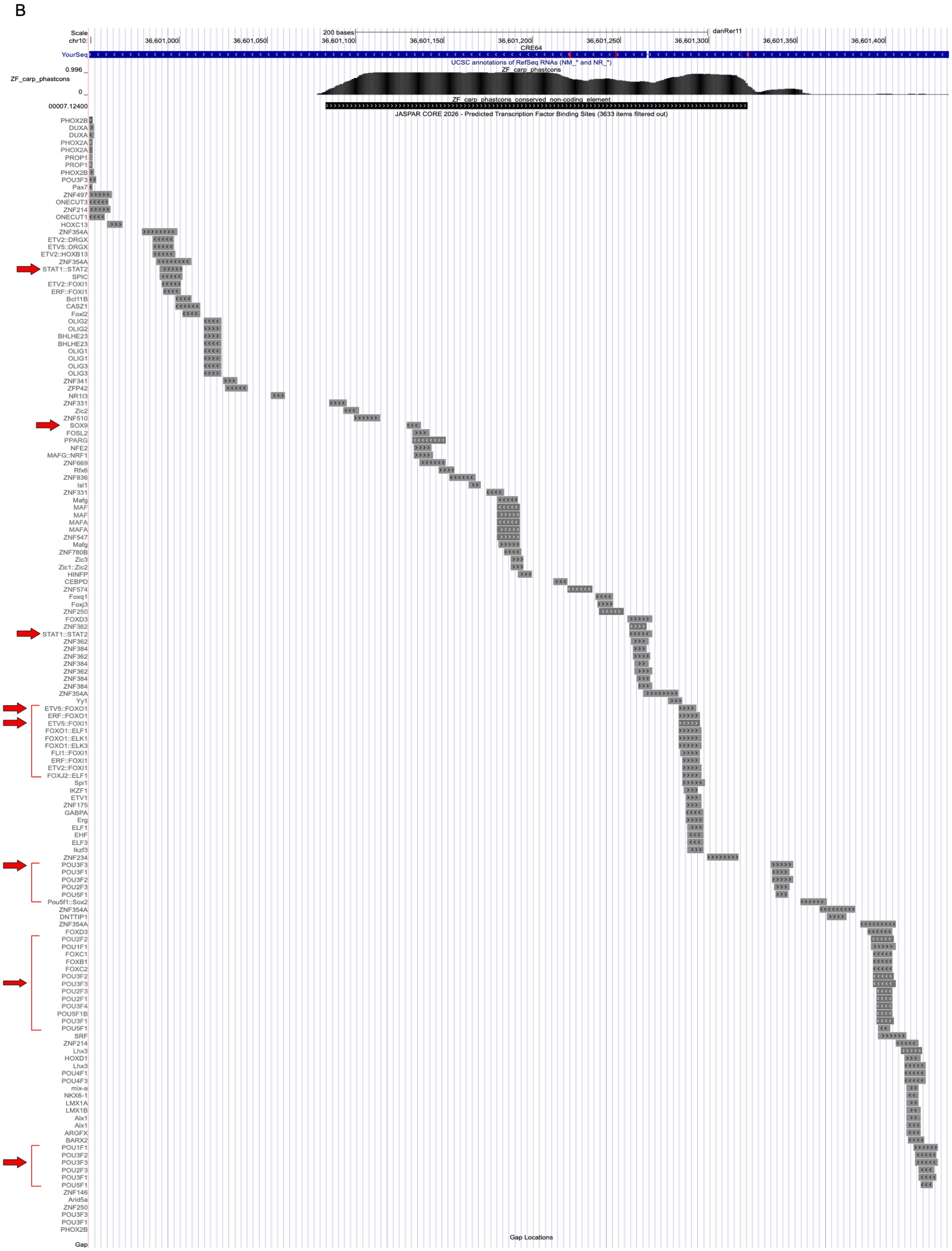


**Supplemental Fig. S13. Predicted transcription factor binding sites at CRE63 and CRE64 of the *slc1a3b* locus.** (A) Zoomed-in view of the min200–400 interval within CRE63. Predicted TF binding sites identified using JASPAR (CORE 2026) are shown, including TEAD and STAT family motifs, which coincide with the conserved sequence block (red rectangles). Red arrows point to predicted TEAD1 and STAT3 binding sites. (B) Zoomed-in view of the conserved region within CRE64. Predicted transcription factor binding sites identified with JASPAR (CORE 2026) are shown, including SOX9, ETV5, POU3F3, and STAT1::2 motifs (red arrows). PhastCons conservation scores across teleost species (ZF-carp alignment) are shown with conserved non-coding elements highlighted.

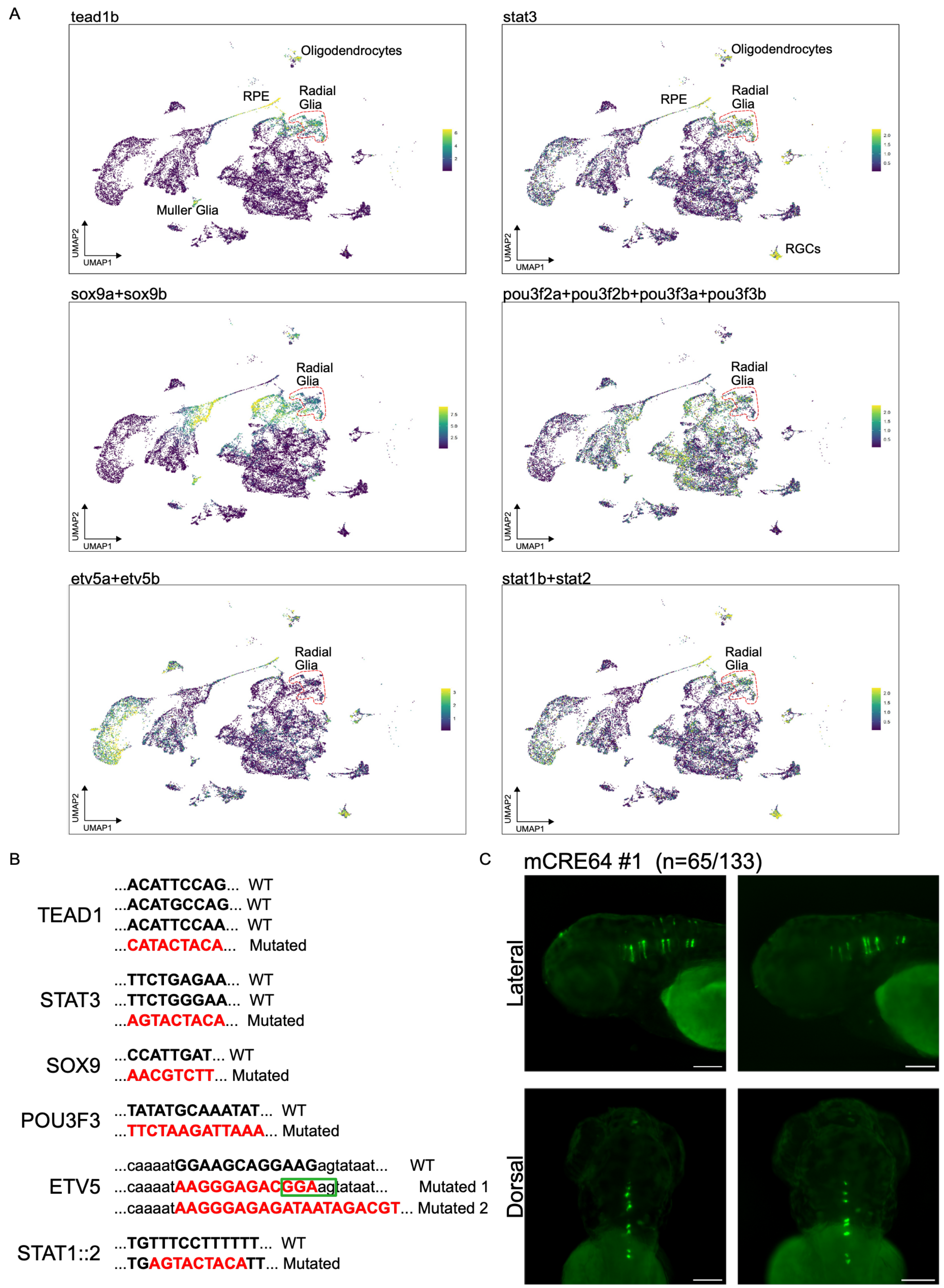


**Supplemental Fig. S14. Motif perturbations of CRE63 and CRE64.** (A) UMAP projections of 3 dpf scATAC-seq data colored by chromVAR motif deviation scores for candidate TF regulators of *slc1a3b* enhancers. Dotted red lines outline radial glia cluster. (B) Wildtype and mutated variants of predicted TF binding motifs within CRE63 and CRE64 enhancers. For ETV5, two mutated sequence variants were tested. Green box highlights a residual GGA core sequence retained in ETV5 Mutated 1 construct. (C) Representative fluorescence images of 3 dpf zebrafish showing mosaic expression of the mutated construct mCRE64 #1. eGFP expression is detected in radial glia but is restricted to the hindbrain region. In contrast, wildtype CRE64 activity is also detected in anterior parts of the brain.

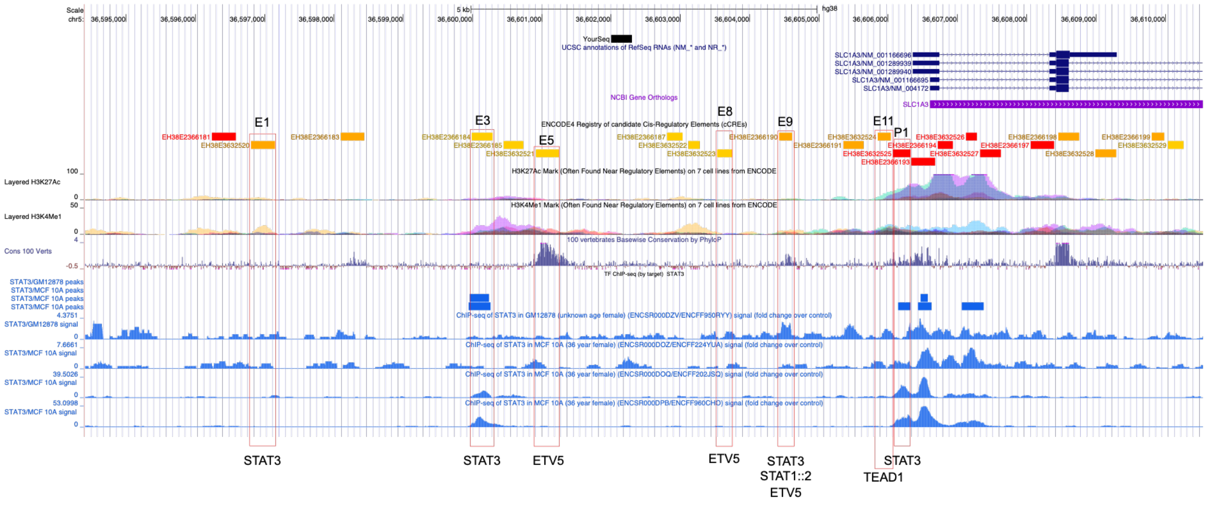


**Supplemental Fig. S15**. **Candidate regulatory elements upstream of the human SLC1A3 locus.** Genome browser view of ~10 kb upstream of the human SLC1A3 locus (hg38) showing ENCODE candidate CREs, histone modification tracks (H3K27ac and H3K4me1), vertebrate conservation (PhyloP, 100 vertebrates), and STAT3 ChIP-seq datasets (GM12878 and MCF10A). Enhancers (E1, E3, E5, E8, E9, E11) containing predicted STAT3, TEAD1, ETV5, and STAT1::2 binding motifs are boxed in red. P1 represents a promoter element.

**Supplemental Tables:**

**Supplemental Table S1.** scATAC-seq cluster sizes and gene activity scores at 3 dpf, 21dpf, and adult stages.

**Supplemental Table S2.** Representative canonical markers used for cell type annotation of 3 dpf, 21 dpf, and adult scATAC-seq datasets along with subtype markers.

**Supplemental Table S3.** Cluster annotations and canonical markers for brain and retina timepoint-integrated scATAC datasets.

**Supplemental Table S4:** Cell and DA peak counts per cluster and timepoint for brain and retina timepoint-integrated objects.

**Supplemental Table S5:** DA peaks identified via pairwise comparisons between timepoints for timepoint-integrated brain and retina clusters.

**Supplemental Table S6:** List of identified peak-to-gene links and GO enrichment results for DA peaks in brain and retina at 3 dpf and adult stages.

**Supplemental Table S7.** chromVAR transcription factor motif enrichment analysis across 3 dpf, 21 dpf, and adult scATAC-seq datasets.

**Supplemental Table S8.** Overlap of enriched transcription factor binding motifs within cell types across 3 dpf, 21 dpf, and adult stages.

**Supplemental Table S9.** Primers and genomic coordinates of candidate cis-regulatory elements tested for enhancer activity in vivo.

**Supplemental Table S10.** List of mfge8a HCR-RNA FISH probe sequences.
